# Omics integration analyses reveal the early evolution of malignancy in breast cancer

**DOI:** 10.1101/2020.04.09.033845

**Authors:** Shamim Sarhadi, Ali Salehzadeh-Yazdi, Mehdi Damaghi, Nosratallah Zarghami, Olaf Wolkenhauer, Hedayatollah Hosseini

## Abstract

**Background:** The majority of cancer evolution studies are done on individual-based approaches that neglect population dynamics necessity for the global picture of cancer evolution in each cancer type. Here, we conducted a population-based study in breast cancer to understand the timing of malignancy evolution and its correlation to the genetic evolution of pathological stages.

**Results:** In an omics integrative approach, we integrated gene expression and genomic aberration data for pre-invasive (DCIS, early-stage) and post-invasive (IDC, late-stage) samples and investigated the evolutionary role of further genetic changes in late stages compared to the early ones. We found that single genetic alterations (SGAs) and copy number alterations (CNAs) conspire together for the fine-tuning of the operating signaling pathways of tumors in *forward* and *backward evolution* manners. The *forward evolution* applies to new genetic changes that boost the efficiency of selected signaling pathways. The *backward evolution*, which we detected for CNAs, is a mean to reverse unwanted SGAs of earlier stages. Analyses of the integrated point mutation and gene expression data show that (i) our proposed fine-tuning concept is also applicable in metastasis, and (ii) metastasis diverges from primary tumor sometimes at the DCIS stage.

**Conclusions:** Our results indicate that malignant potency of breast tumors is constant over pre and post invasive pathological stages. Indeed, further genetic alterations in later stages do not establish *de novo* malignancy routes; however, they serve to fine-tune antecedent signaling pathways.

## Background

Modern cancer biology, which is expanding rapidly in the light of omics approaches, validates many aspects of Darwinian somatic evolution in cancer progression^1^. Such evolutionary logic influences and adds practical benefits to our thinking and actions^2^. However, there are still some cavities in the cancer evolution field, requiring new thinking and approaches to be addressed. For example, there is not clear how the progress of tumor staging could be correlated to the clinical outcomes as the final phenotypic readout of cancer evolution. It is speculated that tumor staging is the leading prognosis indicator and also decline in cancer mortality in diverse cancer types is predominantly the result of screening that inhibits the incidence of larger tumors^3-8^. However, it has been suggested that features of the tumors have more prognostic relevance than the anatomy of tumor^9^. On the top, recent studies in breast and colorectal cancer have shown that disseminated cancer cells from earlier stages are the main founder of metastases^10,11^. Hence, it is still unclear to what extent the evolution of malignancy represents the evolution of genetic alteration and pathological progression of tumors.

The study of the somatic evolution of cancer relies on the view of dynamic genotype-phenotype relationship of cancer cells at different stages of tumor progression^12^. The dynamic view for cancer evolutionary studies are mostly obtained by paired or longitudinal sampling from single individual patients^13,14^. However, this approach is technically complicated and has some limitation by defaults. For example, multi sampling and biopsies are not adequate to cover the whole heterogeneous tumor population^14,15^. Moreover, the depth of sequencing which is used to get the mutational portrait of samples is still below the borders to detect minor sub-clones within the samples^16^. Most importantly, the high impact of genetic drift, genetic background, and neutral evolution introduce undetectable bias in the individual-based evolution studies^17,18^. Recently, Pan-cancer analysis attempts to draw maps of similarities and differences among the genomic and cellular alterations in a population basis across diverse tumor types reflecting some evolutionary hints^19^. Nonetheless, there is still a gap to cover cancer evolution within the population of each cancer type.

Studying the evolution of malignancy in large populations of patients will cover some of the mentioned issues and might pinpoint some new aspects of cancer evolution which is not visible in the individual-based approaches. We are exploiting high amount of available data on different stages of breast cancer as a beneficial to study a universal portrait for the tumor evolution in breast cancer. The most common type of breast cancer is the ductal carcinoma that consists of two main stages including Ductal Carcinoma In Situ (DCIS) and Invasive Ductal Carcinoma (IDC)^20^. The DCIS or stage-0 breast cancer is considered as the non-invasive stage and the immediate precursor of the most invasive breast cancers^20,21^.

In this study, we used an omics integration approach for the gene expression, copy number alteration (CNA), and mutation data for the DCIS, IDC and metastasis (Met) samples. Our results indicated that the evolution of malignant phenotype is mostly forming in the earlier stages of tumor progression. Phylogenetic analyses depicted a global picture of breast cancer evolution that indicated early divergence of metastases at DCIS stage. We discovered that further genetic alterations in the advanced tumors do not generate new paths for the function of tumor but rather serve as fine-tuning effectors for antecedent changes from earlier stages. The fine-tuning happens in two different paths as we called them *forward* and *backward evolution*. In *forward* evolution, cancer cells increase the single genetic alterations (SGAs) burden to increase the efficiency of selected pathways. On the other hand, in the *backward evolution* program, tumor cells exploit CNAs in order to reverse the expression direction of unwanted inherited genetic changes with negative impact on the function of operating pathways. Therefore, we concluded that malignancy fate is determined at very early stages and further genetic changes are the fine tuner of evolutionary path.

## Results

### DCIS and IDC are indistinguishable based on the gene expression profile

General belief in breast cancer progression sees DCIS as the non-invasive and IDC as the invasive state of breast tumors^20,21^. We hypothesized that DCIS can be dissociated from IDC based on their molecular features. Hence, we assessed the uniqueness of normal from DCIS and IDC samples and DCIS from IDC using gene expression analyses. Accordingly, we used 10 datasets including 430 DCIS, IDC, and normal samples, profiled by three different platforms (Fig S1a). We used microarray data due to the availability of sufficient gene expression data for DCIS and IDC samples with a normal matched datasets. Then, we performed statistical analysis and class comparison test for identification of differentially expressed genes (DEG) that resulted in 4275, 5000, and 1098 DEGs with a combined *p*-value < 0.05 for DEG1 (DCIS vs Normal), DEG2 (IDC vs Normal), and DEG3 (DCIS vs IDC), respectively (SI File 1, Fig S1b). In order to be sure that our results are not biased because of the faults of microarray technology, we compared DEG2 with an analogous DEG from the largest available RNA-sequencing dataset in breast cancer^22^ and found significant similarities between these two data types (p<0.0001, Exact hypergeometric probability test; Fig S1c).

After DEG identification, we used principal component analysis (PCA) and hierarchical clustering to investigate if the gene expression profiles separate the DCIS and IDC samples. Indeed, we did not find a clear separation between these samples (Fig 1a, and b). To avoid the bias of incompatibility noise in our analyses, we used COMBAT in the PCA and hierarchical clustering methods to remove the batch noises. Applying PCA and hierarchical clustering method after COMBAT could dissociate DCIS and IDC samples from normal samples. However, DCIS and IDC samples could not be dissociated even after applying COMBAT (Fig 1a, and b). These results indicate that DCIS is not distinguishable from IDC using gene expression clustering approaches.

**Figure 1.**
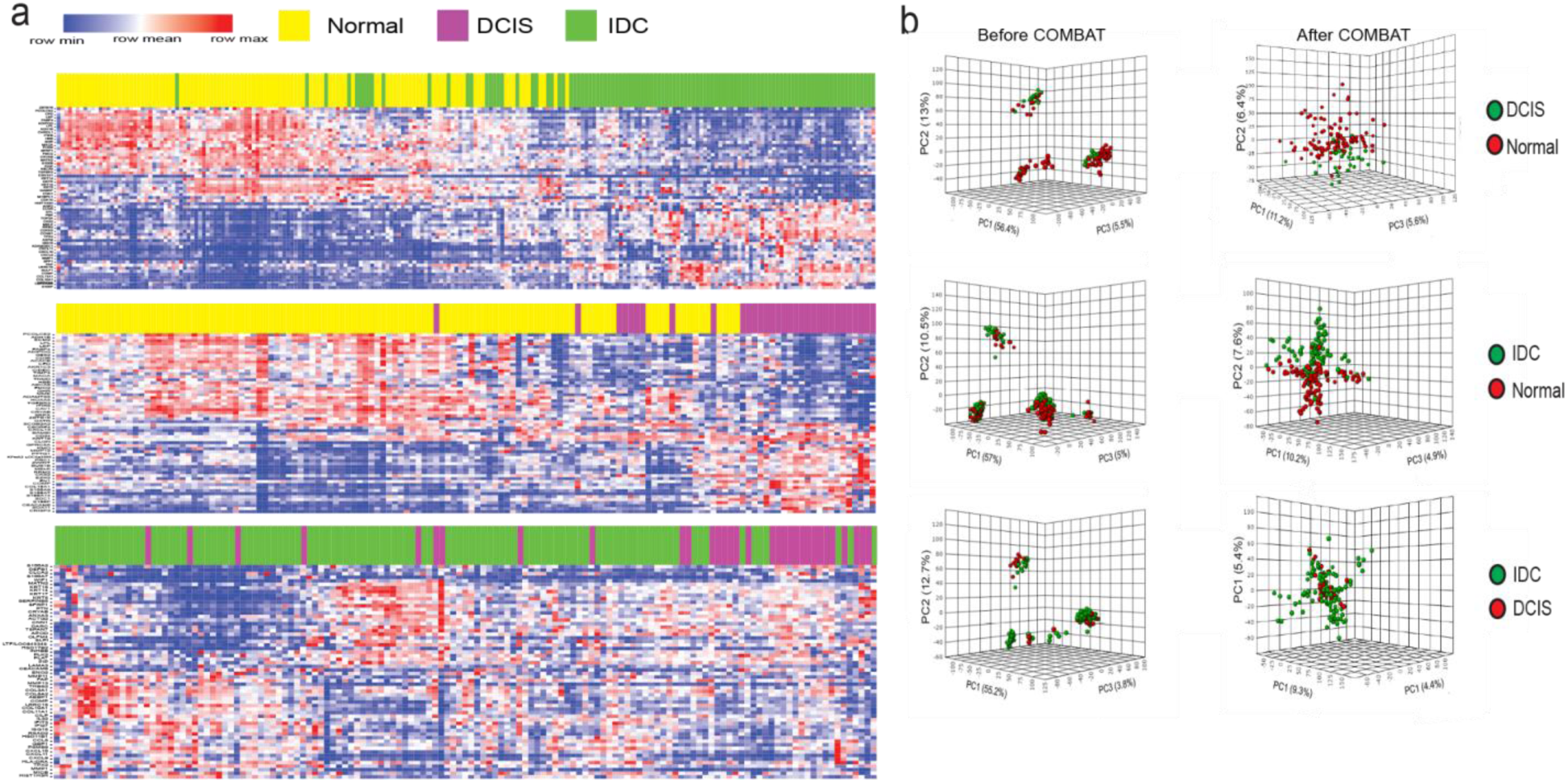
Unsupervised hierarchical clustering and PCA. **a**) Two-dimensional heatmaps show 60 DEGs based on the log fold changes. Clustering was done after applying COMBAT method. **b**) PCA analyses before (left) and after COMBAT (right). PCA distinguishes DCIS and IDC from normal samples based on their gene expression profile but it failed to distinguish DCIS from IDC samples.

### Stage-dependent gene expression analyses do not generate a malignancy-predictive feature

Here, we applied further advanced approaches to distinguish DCIS and IDC based on their gene expression profiles. To do this, we used machine learning (ML) methods to define a minimal number of genes as representatives of DCIS and IDC. We optimized four different ML methods (weighted Voting X Validation, KNN, GBM, and Random Forest) and they were successfully generating distinctive (ROC/AUC>0.5) features to sort out DCIS from IDC samples (Fig 2a). In order to weight the accuracy of selected gene features, we removed all genes of the first selected features and re-ran all ML models to get new distinctive features. Notably, all ML methods were able to reconstruct new distinctive features even after excluding previous selected gene features from the data source (Fig 2b and Fig S2a). Next, we hypothesized that the first selected gene features may have a stronger biological correlation to the clinical outcomes (e.g., survival) in patients as compared to the gene features in the next rounds (Fig 2c and Fig S2b). Our analyses revealed that the discriminative power for the selected features of our chosen ML methods (ROC/AUC) does not indicate their biological relevance. Indeed, features with non-significant discriminative power (ROC/AUC <0.5) were also strongly correlated to biological outcomes (survival analyses Fig 2c and Fig S2b). Inspecting the survival analyses and selected features by ML methods, we realized that the strength of the hazard ratio (HR) and concordance index (CI) is strongly correlated to the number of genes in the selected features (p<0.0001, Pearson correlation test; Fig 2d). Furthermore, we could not find a significant consistency between the direction of expressed genes in the source of selected feature gene (DCIS or IDC) and the poor and good prognosis samples in the survival analyses (Fig S2c). These results suggest the dissociation power of the discriminative gene features of DCIS versus IDC is not correlated to the clinical outcomes and malignancy of breast tumors.

**Figure 2.**
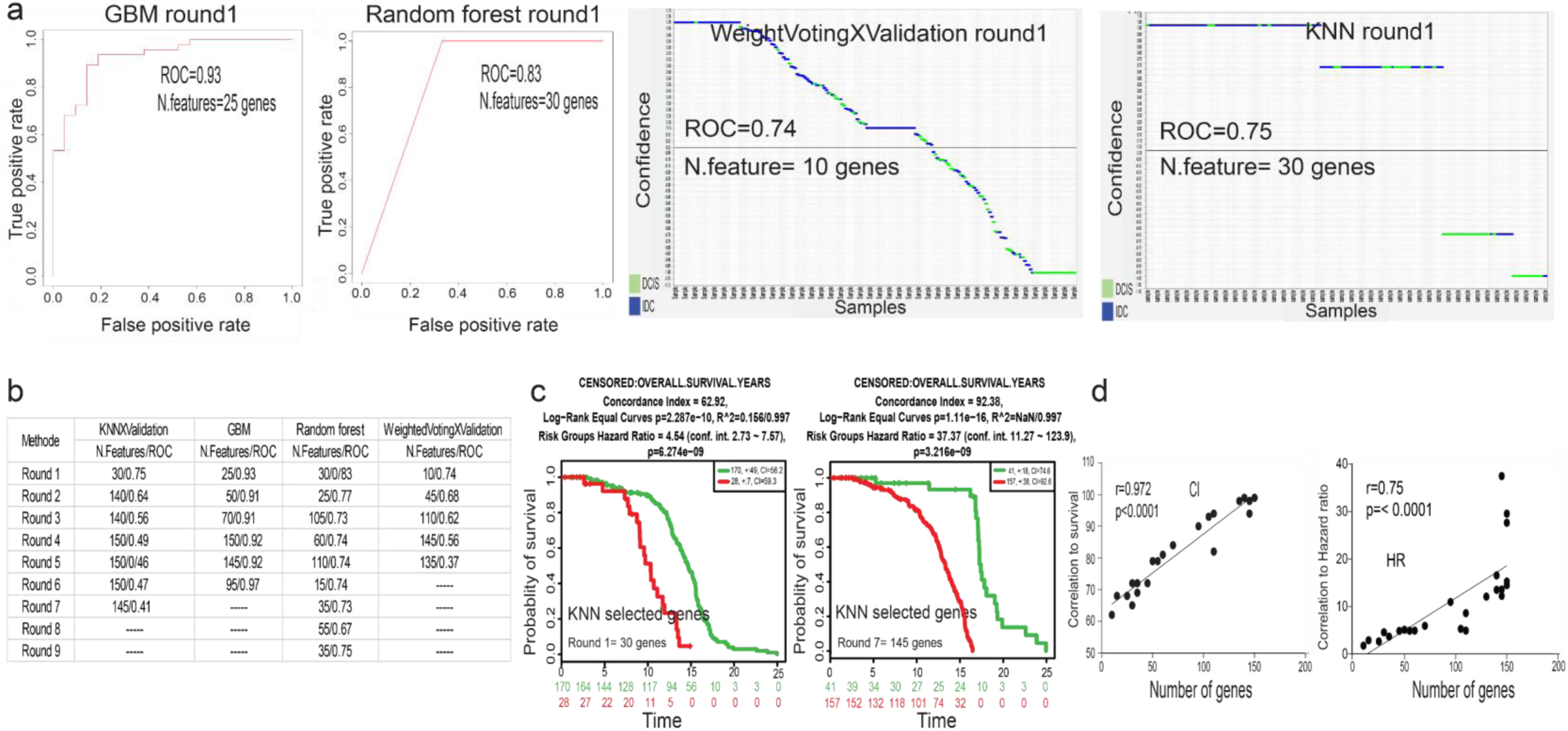
Machine learning predictive models and survival analyses. **a**) ROC plots and plots obtained by prediction results viewer for the first round of feature selection by four different machine learning (ML) models. Separation of DCIS and IDC samples were identified by both models. **b**) ROC results of first nine rounds of feature selections. After each round of feature selection, we excluded this gene list feature from DEG source and repeated for a new feature selection. Note that we did not continue feature generation if ROC/AUC was very constant or dropped below 0.5. **c**) Kaplan-Meier survival analyses of the first and the last rounds of gene feature selection optimized by KNN model. Note that the last feature with ROC below 0.5 (panel b) also strongly correlated with survival. All log rank tests are significant (*p*<0.05; *p*-value generated by SurvExpress online server) **d**) Correlation of concordance Index (CI; left) and hazard ration (HR; right) with the number of genes that were obtained from optimization of each round of models (Pearson correlation test).

### Stage-dependent CNA and SGA integrative analyses do not generate malignancy-predictive features

The aggravation of chromosomal aberration is a general aspect of tumor progression in the evolutionary path toward advanced stages^23^. We speculated that the integration of CNA (Copy Number Alteration) and SGA (Single Gene Alteration, extracted from DEGs) would help to find a better discriminative feature for sorting out DCIS from IDC samples. Indeed, we hypothesized that the integration of SGAs and CNAs would generate better discriminative features to dissociate DCIS and IDC in correlation with malignancy outcomes (e.g. survival analyses). To test this hypothesis, we extracted the CNA map of DCIS and IDC samples (Progenetix, database; version 2017; 76 DCIS and 1,736 IDC samples; Fig 3a; SI File 2). We considered all IDC had experienced a DCIS stage during tumor progression. Thus, we generated a combined map of the CNAs of DCIS and IDC to depict an evolutionary view on the CNA map for breast tumors (Fig 3b). This map generated 25 prototypes including, five prototypes were specific for DCIS and three prototypes were specific for IDC. There were two complex prototypes without a clear gain and loss patterns in correlation with DCIS or IDC stage (see methods and SI File 2 and 3). We also found 15 shared prototypes between DCIS and IDC. On the other hand, not all SGAs were *de novo* generated in IDC. Almost half of the SGAs (508 out of 1,098 for DEG3, IDC vs DCIS) originated from DCIS, as we call them *common genes*, and their changes became stronger in IDC (Fig 3c, Fig S3a and SI File 1). The rest of the SGAs which we call disparate genes including SGAs generated exclusively in IDC stage (Fig 3c, Fig S3a and SI File 1). Next, we distributed the SGAs in CNAs and ranked them based on log fold changes (LFC). Then, we tested if any of these gene sets (common/disparate, within/out CNAs) has a better CI to the survival outcomes. For this analysis, we selected features of 10 genes (arbitrary number) and added the number of genes by 10 more genes for the next features. We performed these analyses for the fold changed ranked gene lists and for the random list. Interestingly, the CI of all groups of SGAs was correlated to the number of genes in each feature and not their CNA related category (Fig 3d and Fig S3b). We further assessed whether SGAs-enriched parts of chromosomal regions within CNAs (regions with as many as SGAs with not more than 5Mb distance to the next SGAs; SI File 3) might have a stronger correlation to malignancy outcomes compared to other chromosomal regions. Our results recapitulate previous findings that the number of genes defines the strength of CI and not the specific region (*p*=0.008, Pearson correlation test; Fig 3e). Furthermore, we could not find a significant consistency between the direction of expressed genes in the source of selected feature genes (IDC) and the poor and good prognosis samples in the survival analyses (Fig S3c). Finally, we tested several well-known gene signatures which are used in the clinic and found that number of genes in these signatures strongly correlated to their CI power (Fig S3d). Altogether, these results suggest that the increase of genetic alterations in IDC as compared to DCIS does not necessary increase the malignant properties of tumors.

**Figure 3.**
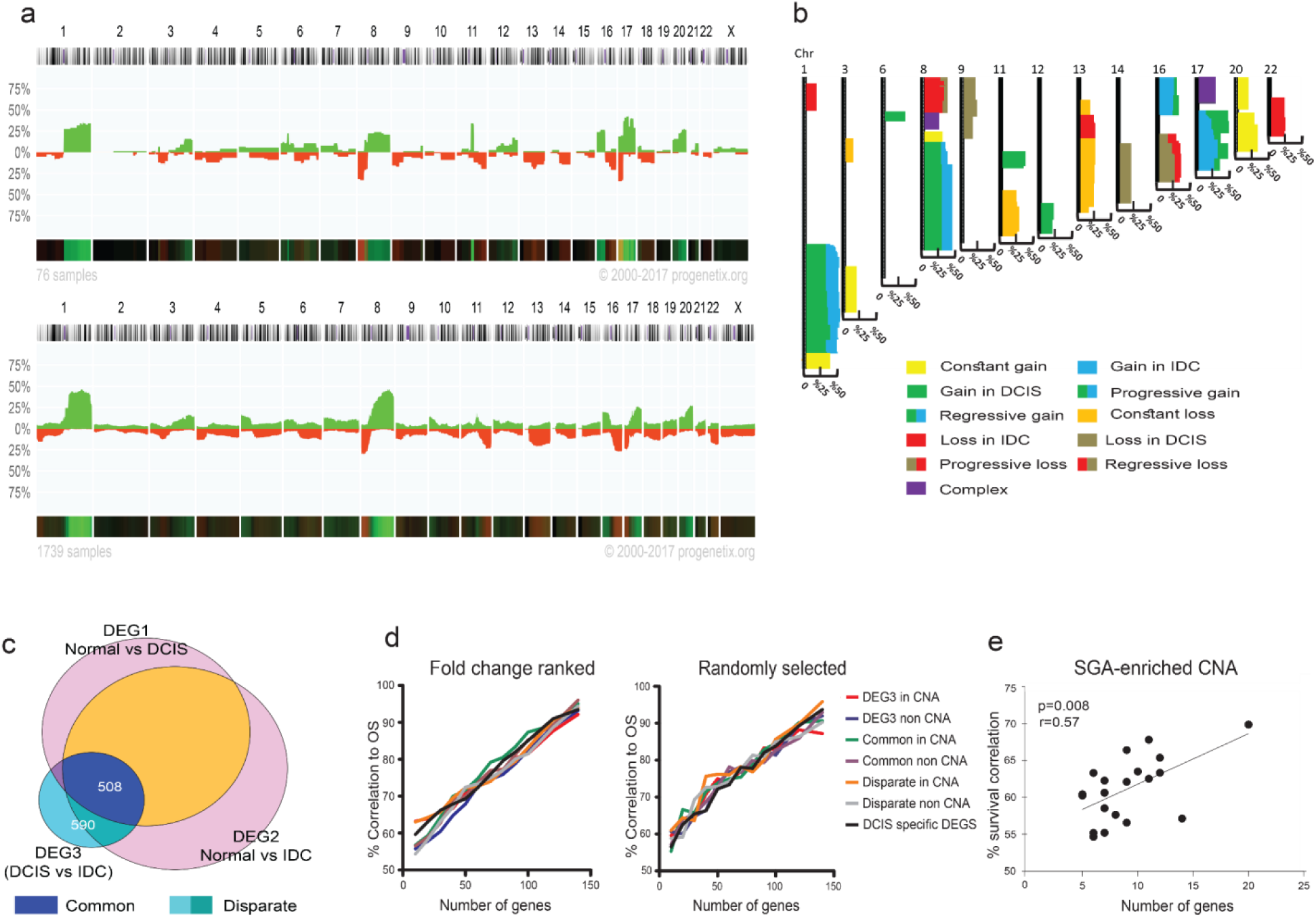
Integration of copy number alteration (CNA) and single gene alteration (SGA) of DCIS and IDC. **a**) CNA map of DCIS (up) and IDC (down) obtained from Progenetix database (DCIS=76, IDC=1739). The *y-*axis depicts the percentage of samples with aberrations (green, gain; red, loss) for each chromosomal region. **b**) Map of CNA prototypes (See also SupFile3). **c**) A Venn diagram scheme of common and disparate genes. **d**) Correlation of number of genes that selected based on ranked fold changes (left) or randomly (right) to overall survival. Genes are selected from common, disparate, DCIS specific and DEG3 (IDC vs DCIS) within and out of CNAs. **e**) Correlation of number of genes in SGA-enriched CNA to survival. *p*-value in panel e calculated by Pearson correlation test.

### Genetic changes in earlier tumor stages are more functional than in later stages

The constant malignant properties of tumors in pre-malignant stages raise a question about the role of accumulation of genetic alterations in the later stages. To address this question, we firstly need to understand the functions and the possible interactions between earlier and later events during tumor evolution. Thereby, we assumed each CNA prototype as an evolutionary unit (Fig 3b) where we can assess the correlation between the direction of CNA and SGA changes during the progression of tumors from DCIS to IDC. The IDC has two types of gene expression profiles; common as originated from DCIS and disparate, which is acquired specifically in IDC (see Fig 3c and SI File1). In addition, there are different types of CNAs for IDC and DCIS; DCIS-specific which only exist in DCIS, progressive CNAs which are inherited from DCIS and progressed further in IDC, IDC-specific which are acquired specifically in IDC, and constant CNAs which are constant changes (gain or loss) in DCIS and IDC. First, we calculated the density of common and disparate up- and down-regulated genes in their CNA prototype (SGA-density is the number of SGA/Mb; SI File 4). Then, we specifically tested whether the expression direction of common and disparate SGA density is similar to the direction of CNAs in each stage (Fig S4a). Our results revealed a positive correlation between the expression-direction of SGAs and the direction (gain or loss) of CNAs in each tumor stage (p=0.008 for common genes in Constant-CNAs and DCIS-CNAs, p=0.06 for disparate genes in Constant-CNAs IDC-CNAs, Wilcoxon matched-pair signed rank test; Fig 4a-c and Fig S4a-b). Then, we sought a serial relation between common and disparate SGAs and their correlation to the direction of CNAs. Since the direction of changes of SGAs and CNAs in the stage-dependent analyses were harmonic and DCIS and IDC had indistinguishable gene expression profiles, we speculated a positive correlation between the direction of IDC-CNAs and the direction of changes for common and disparate SGAs together. Here, we hypothesized a *forward evolution* model in which the disparate SGAs increase the density of SGAs in the direction (up- or down-regulation) of existing common SGAs in the IDC-specific and IDC-progressive CNAs (Fig 4d and S4c). Surprisingly, we could not confirm the *forward evolution* model for the consequential common and disparate SGAs in the CNAs related to IDC (Fig 4d-f). Subsequently, we analyzed the functional interaction of common and disparate SGAs using protein-protein interaction networks (PPINs). Because of the last results, we separated SGAs to two categories; SGAs within and out of CNAs. Interestingly, we found that common SGAs were the most interactive in the specific PPINs analyses, because of the higher average node degree (Fig 4g). Notably, SGAs within CNAs were poorly connected in the networks. In addition, gene set enrichment analyses (GSEA) revealed the enrichment of more functional pathways for the common gene sets compared to disparate genes and SGAs in the CNAs (Fig S4d).

**Figure 4.**
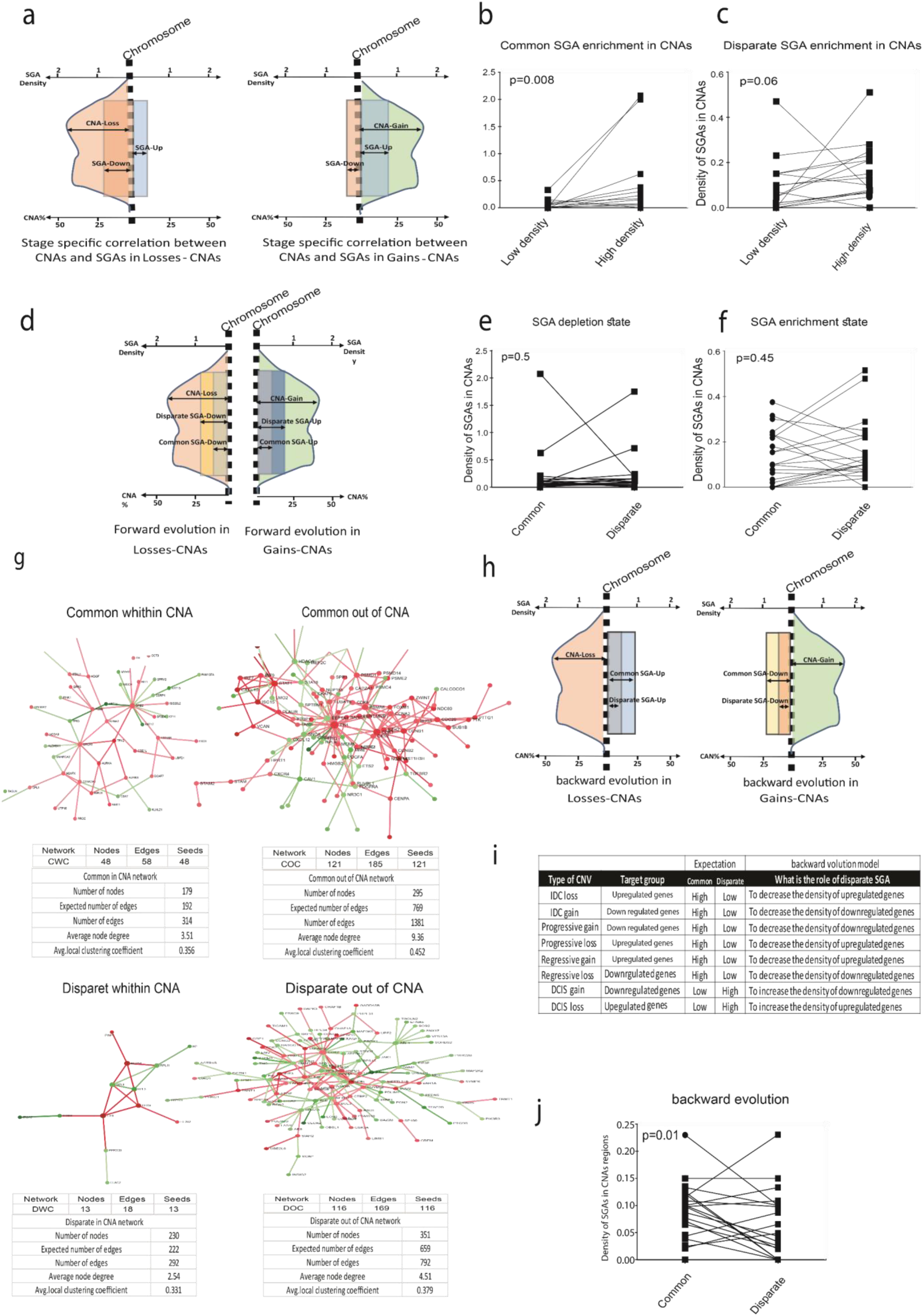
Evolutionary correlation models of SGA and CNA. **a**) This scheme depicts the expectation of stage specific correlation between the direction of CNAs and SGAs. It is expected that the density of up-regulated-SGAs be higher than down-regulated-SGAs in gain-CNAs (right) and the density of down-regulated-SGAs be higher than up-regulated-SGAs in loss-CNAs (left). **b**) Common SGAs are correlated with the direction of CNA prototypes of DCIS. Density of down-regulated genes is high in loss-CNA and density of up-regulated genes is high in gain-CNAs (See Fig S4a). **c**) Disparate SGAs are correlated with the direction of CNAs prototypes of IDC. The density of down-regulated genes is high in loss-CNAs and density of up-regulated genes is high in gain-CNAs (See Fig S4 b). **d**) This scheme depicts the expectation of *forward evolution* for the common and disparate SGAs in IDC-CNAs. It is expected that the density of disparate upregulated-SGAs be higher than common upregulated-SGAs in gain-CNAs (right) and the density of disparate downregulated-SGAs be higher than common downregulated-SGAs in loss-CNAs (left; see also Fig S4c). **e** and **f**) The *forward evolution* model is not confirmed neither for the SGA-depletion state (e) nor SGA-enrichment state (f; see Fig 4c). **g**) Specific PPIN of common SGAs within CNAs (CWC), common out of CNAs (COC), disparate within CNAs (DWC) and disparate out of CNAs (DOC). Networks were constructed by mapping gene sets to human PPIN database. Green and red nodes represent down- and up-regulated genes, respectively. PPINs statistics is calculated by STRING. **h**) This scheme depicts the expectation of *backward evolution* for the common and disparate SGAs in IDC-CNAs. It is expected that *backward evolution* decreases the density of upregulated-SGAs in loss-CNAs (left) and the density of downregulated-SGAs in gain-CNAs (right). **i**) Scheme table presents a *backward evolution* concept (Numbers for this prediction presented in Fig S4e). Note that we evaluated all SGAs (common and disparate) in the IDC-CNAs or progressive CNAs prototypes as the final evolved version of CNAs. **j**) Evaluation of *backward evolution. p*-values in panels b, c, e, f, and j calculated by Wilcoxon matched-pair signed rank test.

Thus far, our results suggested that progression of CNAs in later stages doesn’t increase the density of SGAs in each given CNA. Surprisingly, further genetic alterations which are specific for later stage don’t generate further signaling pathways in the tumor that is falsifying a forward evolution model for the role of extra genetic alterations in later stages. Hence, we speculated two scenarios to explain the role of further genetic alterations in the later stages. The first scenario is to consider a neutral evolution model where further genetic alterations randomly accumulated in tumors. This scenario can be true but difficult to test it. In a substitute scenario, we hypothesized a *backward evolution* in the later stages that serves to reverse the expression of unwanted genes inherited from earlier stages. In a *backward evolutionary* model, we supposed that the direction of CNAs in the later stages function in the reverse direction of SGAs from earlier stages within that given region (Fig 4h-i). For example, a *forward evolution* model supposed an additive impact where an IDC-CNA gain, increases the density of up-regulated SGAs (which failed; Fig 4d-f), however, the *backward evolution* model posits that an IDC-CNA gain, is selected to decrease the density of down-regulated SGAs in that given chromosomal region which inherited from DCIS stage (Fig 4i). Our results corroborated this hypothesis where we found that later CNAs reversely change the density of common SGAs of those regions (p=0.01, Wilcoxon matched-pair signed rank test; Fig 4j; Fig S4e). This hypothesis is further supported by analyzing of CNAs being constant (the percentage of gain or loss is constant) through DCIS to IDC stages. In these constant CNAs, the density of common and disparate SGA did not change over the stages; neither in the *forward* nor in the *backward* models (Fig S4f-h). Therefore, we concluded that CNAs could serve as *backward evolutionary* tools in later stages to reverse the expression of some of the inherited SGAs from earlier stages.

### *Forward* and *backward evolution* models are exploited for the fine-tuning of operating pathways during tumor progression

To better understand the role of early and late genetic alterations during tumor progression we performed more functional analyses. Since tumor progression is an unceasing series of events and existing genetic alterations in later stages are accumulated events during progression from early to late stage, we studied the function of early genetic alterations (e.g. common SGAs) as a part of whole running program in the late tumors. Therefore, we performed PPIN analyses for common SGAs and all DEG3 (all common and disparate SGAs) independent of their chromosomal loci. Surprisingly, we found the higher connectivity of network suggests more intricate biological functions for the protein network of all DEG3 compared to the common SGAs (Fig S5a). Then, we performed GSEA for both networks and found that the common SGA profile was mostly related to cell proliferation pathways. Interestingly, all of these proliferation related pathways were enriched in the analyses of DEG3 profile that indicates the constant role of selected pathways during tumor progression (Fig S5b and SI File 6).

Then we investigated how further genetic alterations may affect the function of selected pathways from earlier stages. Closer inspection of PPINs suggested that high-interactive nodes in the common PPINs are more homogenous in their connections compared to the PPINs of DEG3 (similar color, which means similar direction of expression; Fig 5a and Fig S5c). Thus, we assessed the homogeneity and heterogeneity of high-interactive protein nodes in the PPINs. For this purpose, we analyzed all nodes with ≥ 4 connections and considered them as homogeneous nodes if ≥ 75% of connections showed similar direction of expression (up-/down-regulation). Our analyses revealed that the homogeneity of high-interacting protein nodes in common genes as originated from DCIS stages was 2-3 times larger than at later stages (p<0.0001, chi-square test; Fig 5b and sup SI File 5). These results, along with the information from GSEAs (Fig S5b), suggest that earlier operating pathways during tumor progression stay constant, however, the relation of involved components become more complex.

**Figure 5.**
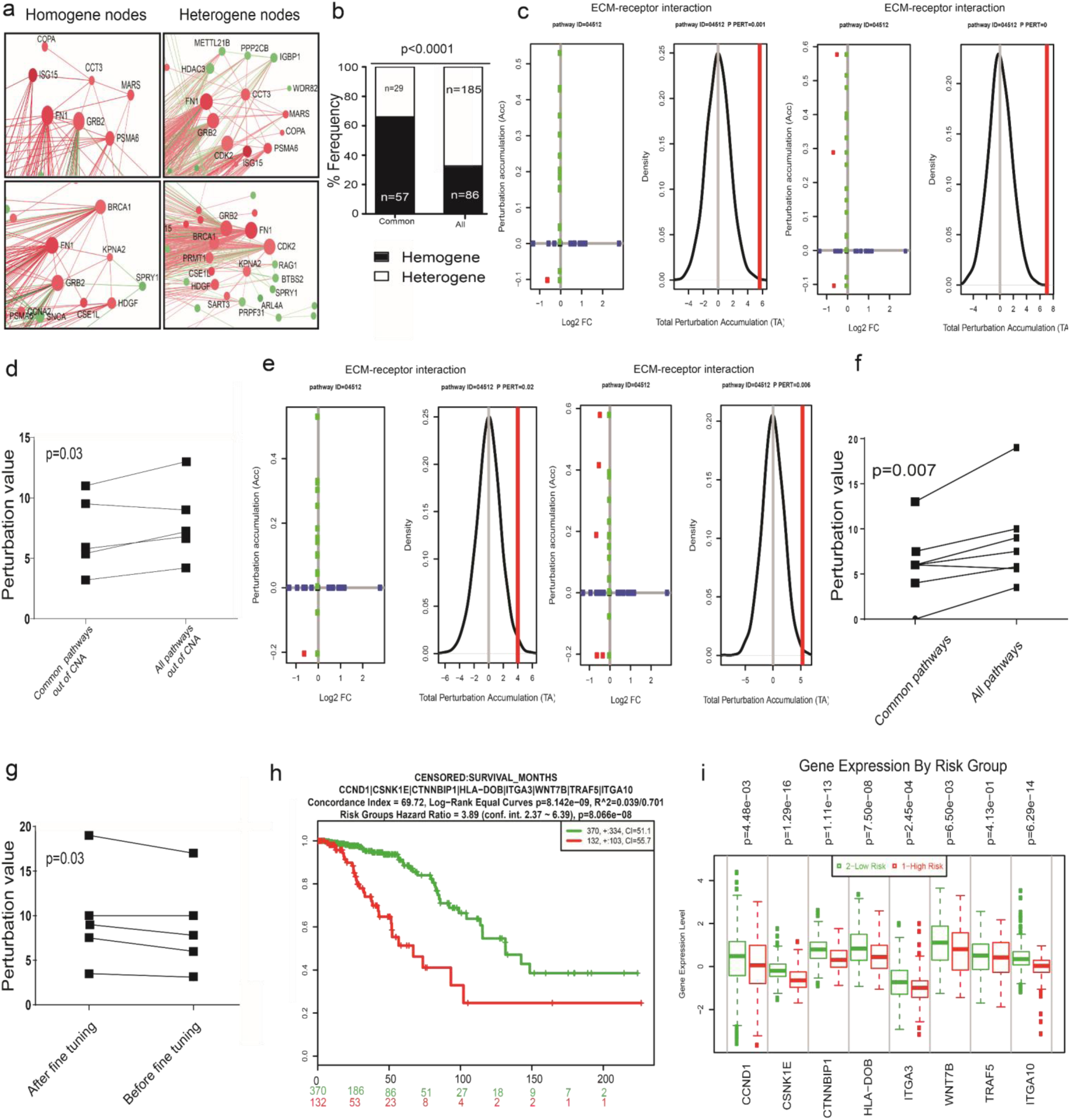
*Forward* and *backward evolution* and *fine-tuning* concept. **a**) Examples of the direction of changes in homogene and heterogene nodes. **b**) Homogeneity and heterogeneity statistical analysis for common and all (all DEG3) networks. **c**) Examples of signaling pathway impact analyses (SPIA) for analyses of common SGAs and all DEG3 (common+disparate) out of CNAs (See Supplementary File 7). In this plot, the horizontal axis represents the *p*-value (minus log of) corresponding to the probability of obtaining at least the observed number of genes (NDE) on the given pathway randomly. The vertical axis represents the *p*-value (minus log of) corresponding to the probability of obtaining the observed total accumulation (tA) or more extreme on the given pathway randomly. Unchanged genes are assigned 0 log2 fold-change. The null distribution of the net accumulated perturbations is also given (right panel). The observed net accumulation tA with the real data is shown as a red vertical line (see ref 27). **d**) Evaluation of perturbation between common SGAs and all DEG3-SGAs out of CNAs (See Supplementary File 7). Note that the numbers presents the pure changes of perturbation compared to the supplementary file 7 which shows negative perturbation values for the inhibition of pathways and positive perturbation value for the activation of pathways. **e**) Examples of SPIA analyses of common SGAs and all DEG3 (See Supplementary File 7). **f**) Evaluation of perturbation between common SGAs and all DEG3-SGAs. **g**) Evaluation of perturbation between enriched pathways for all DEG3-SGAs before adding eight *backward evolution* genes and after adding *backward evolution* genes. **h** and **i**) Survival analyses for eight *backward evolution* genes (h). The direction of backward evolved genes is in the direction of good prognosis samples (i). Evaluation *p*-values in panels d, f, and g calculated by paired student t-test and in panel b, k and l by chi-square test.

To investigate the functionality of the acquired complexity, we expanded our analyses by using signaling pathway impact analyses (SPIA) to track the function of changes in the enriched pathways. The biggest advantages of SPIA is its ability to quantify the direction of changes by integrating the effect of topological positioning, gene expression values, and the impact of players interaction (inhibition or activation) in that given enriched pathway^24^. Thereby, we performed SPIA for (1) common SGAs out of CNAs as earlier event in comparison with all SGAs of DEG3 out of CNAs, and (2) all common SGAs as compared to all DEG3 to understand the functional impact of SGAs within the CNAs in the enriched pathways. Our analyses for the SGAs (only SGAs out of CNAs) show that only five pathways have significant perturbation for the common profile (Fig 5c, Supplementary file 7). Next, we found perturbation of enriched pathways for common SGAs increases when we tested all SGAs (only SGAs out of CNAs; Fig 5d; p=0.03 student paired t-test) that indicate SGAs increase the function of selected pathways in a *forward evolution* manner. Similarly, we performed SPIA for all SGAs of common genes compared to all DEG3 (all SGAs within and out of CNAs). Interestingly, perturbation was detected for only seven pathways (Fig 5e and Supplementary file 7). Consistently, the perturbation values increased in the same enriched pathway in all DEG3 compared to the common profile (Fig 5f; p=0.007 student paired t-test). These results indicate that further genetic alterations in later stages do not establish new paths for the operating pathways of tumor cells, but rather fine tune and increase the function of descendant pathways, which can be explained by *forward evolution* model.

In the next step, we functionally assessed the existence of *backward evolution* over tumor progression. In the previous section, we showed that CNAs in later stages serve to decrease the load of SGAs inherited from earlier stages in these regions where their function was left unaddressed (see previous section and Fig4-j). Here, we hypothesized that reverting of SGAs by CNAs in the later stages increases the function of selected pathways in later stages. In this order, we sorted all genes from DEG1 (DCIS vs. Normal; those are prone for reversing by CNA in later stages) which are existed in the seven enriched pathways with significant perturbation from SPIAs results of DEG3 (Fig 5f and Supplementary file 7). We found eight DCIS-upregulated genes in the CNAs regions (in six out of the seven enriched pathways) with a supposed *backward evolution* function (their expression reversed by CNA in later stage; CNA regions are listed in Fig S4e) and added them to the DEG3 along with their original gene expression value from DCIS and ran SPIA. Remarkably, perturbation of these six pathways significantly decreased upon adding of the mentioned eight genes from DCIS (Fig 5g; p=0.03 student paired t-test; Fig S5d; Supplementary file 7). We further tested these eight genes for the correlation with the survival of patients with invasive breast carcinoma (TCGA data deposited in Survexpress) which showed significant prognostic value for dissociation of the poor and good prognosis samples. Interestingly, all of these genes have similar (higher) expression to their DCIS source in the good prognosis samples compared to poor prognosis ones (Fig 5h-i).

Altogether, these results indicate that SGAs in advanced stages, which are inherited from earlier stages, play a prominent role in the construction of functional signaling pathways. Those SGAs, which are acquired in later stages, serve as the fine tuner for the function of SGAs which inherited from earlier stages. These results further suggest that CNAs have no driving role in the evolution of functional signaling pathways of tumors and they are rather exploited tools by tumors for the fine tuning in a *forward* or *backward evolution* models.

### Metastasis diverges from primary tumor at early stages

In cancer, metastasis is the demonstration of malignant phenotype evolution of primary tumors. Therefore, we tested the main aspects of our findings in metastatic (Mets) samples. For these analyses, we first generated a DEG for metastases as well as a common DEG list overlap between Met-DEG and DCIS-DEG profile (common-Met, Fig S6a and SI File 8). Then we ran the SPIA and generated pathways with significant perturbation. We identified 24 pathways for Met-DEG and 21 pathways for common-Met with significant perturbation values whereas 19 of them were shared between two analyses (SI File 9). Consistent with our previous results for IDC (fig 5c-f), fine-tuning was also detected between common-Met and Met-DEG pathways with significant perturbation (p=0.001, Student paired t-test, Fig 6a). Further, we ran SPIA for the DCIS-DEG list and compared perturbed pathways between DCIS, IDC, and Met. Interestingly, we found that metastasis-perturbed pathways are similarly shared between DCIS and IDC (Fisher Exact Probability test, Fig 6b-d). However, the number of perturbed pathways was significantly higher in metastases samples (Fisher Exact Probability Test, Fig 6b and e). Next, we ran a modified version of DawnRank^25^ analysis to identify driver genes of each stage (Fig 6f, SI File 10). Our analyses revealed that IDC has more shared driver genes with DCIS as compared to Mets and interestingly, number of shared driver genes of Mets is similar between Met vs DCIS and Met vs IDC (Chi square test, Fig g and h and Fig S6b, SI File 10). Further, Met and IDC have significantly higher number of unique driver genes compared to DCIS implying a divergence from DCIS and further independent evolution (Fig 6j). Our findings which showed i) Met has more unique perturbed pathways compared to DCIS and IDC (Fig 6e), and ii) Met has more unique driver genes compared to DCIS (Fig 6e), suggest that Met has probably more time for an independent evolution as speculated to be the result of a neutral evolution in the absence of selection pressure^18,26^. Thereby, we hypothesized that phylogenetic analyses would show the early divergence of Met at the time of DCIS. Firstly, based on all similarities between perturbed pathways and driver genes (Fig 6d, e, h, and i), we drew a conceptual phylogenetic tree that demonstrated an early divergence of metastasis at DCIS stage. Then, we drew an accurate phylogenetic tree using mutated genes in the shared driver gene profile between DCIS, IDC and Met (Fig 6k) that consistently generated similar picture to the conceptual phylogeny analyses (Fig 6l). Altogether, these results indicate that the evolution of malignant phenotype in breast cancer is much faster than the evolution of pathological stages.

**Figure 6.**
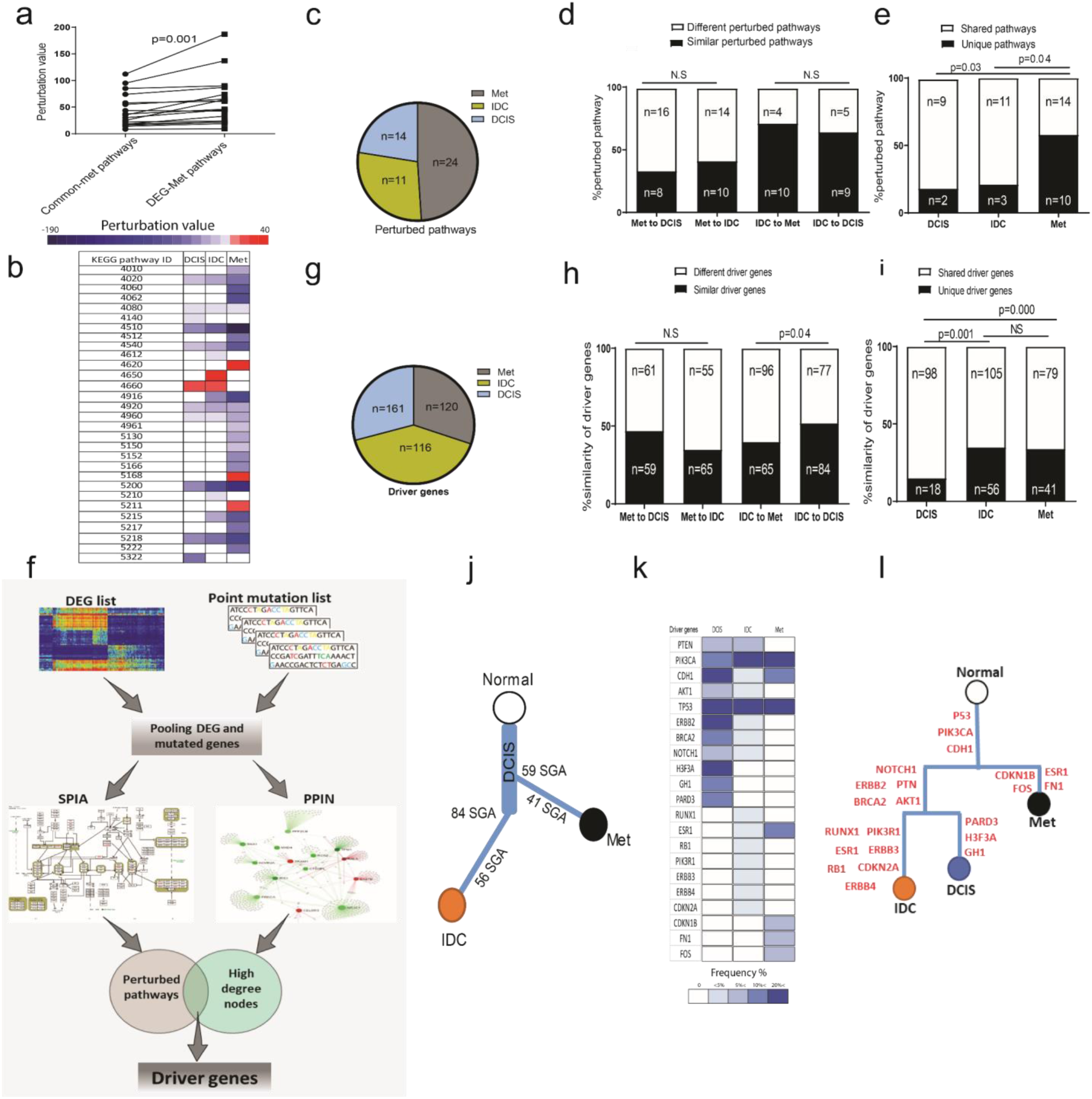
Evolution of metastasis and divergence time from primary tumor. **a**) Evaluation of perturbation between common-Met SGAs and all DEG-Met SGAs (SI File 9). **b**) Heatmap of perturbed pathways obtained by SPIA analyses for DCIS, IDC, and Met (SI File 9). **c**) Pie chart presents number of perturbed pathways in each group. **d**) Similarity of perturbed pathways is shown between DCS, IDC and Met groups (SI File 9). **e**) Comparison shows a number of unique and shared perturbed pathways for each group. The shared perturbed pathway is the one that exists at least in two groups. **f**) Graphic demonstration of our modified version of DawnRank for detection of driver genes (see method). **g**) Pie chart presents number of driver genes in each group. **h**) Similarity of driver genes between DCS, IDC and Met groups (SI File 10, see S6b). **i**) Comparison shows a number of unique and shared driver genes for each group. The shared driver gene is the one that exists at least in two groups (see S6b). **j**) Conceptual phylogenetic tree extracted from the concepts in panels d, e, h, and i. **k**) Heatmap of mutated driver genes shared between DCIS, IDC, and Met (SI File 10, see S6b). **l**) Phylogenetic tree constructed based on mutation profile of shared driver genes between DCIS, IDC, and Met, which presented in panel k. Evaluation *p*-values in panel a calculated by paired student t test, in d and e calculated by Fisher exact probability test, and in h and i calculated by chi-Square test.

## Discussion

As opposed to individual-based approaches, we conducted our evolutionary study in a population basis. This is because in the individual-based approaches, i) sampling methods do not cover the entire heterogeneity of tumors^14-16^, ii) individual sampling has the risk of deviation of interpretation by genetic drift and neutral evolution^17,18^, and iii) the final candidates coming out of individual tumor evolution analyses can be biased by the environment or background of that given individual. In this study, we specifically aimed to address whether the evolution of the malignant phenotype of breast cancer is representing the evolution of pathological stages. We have selected ductal carcinoma, as it is the most frequent breast cancer type^20^ and a reasonable number of samples with their matched adjacent normal tissue. Studying DCIS, IDC, and Met allowed generating an overall view of the stepwise Darwinian evolution of breast tumors.

Our results indicate that different gene expression signatures obtained by different approaches may have discriminative power to separate DCIS from IDC samples, however, these gene expression signatures had nothing to do with the malignant properties of these samples. We further found that the increase of genomic alterations in later stages of tumor progression has no distinctive prognostic value. Our results indicated that SGAs, which are inherited from DCIS, play a prominent role in the construction of operating signaling pathways in later stages. Genetic alterations, which were obtained in later stages, have a fine-tuning role for the function of genetic alterations inherited from earlier stages. Indeed, the majority of signaling pathways with significant perturbation value in DCIS also enriched in IDC and Met. However, the same pathways had three main differences in the advanced tumors as compared to their ancestral pathways from early stage. First, the number of genes in the same pathways is higher in later stages compared to the ancestral ones that indicate the involvement of more genomic changes in the later stages. Second, the same pathways are increasingly heterogeneous in the direction of the expression-regulation of their player in later stages. Third, the perturbation of enriched pathways significantly increases in the late stage that indicates fine-tuning evolution of early-developed pathways by further genetic alteration in late stage tumors.

We also found that the density of SGAs in the chromosomal regions with CNAs is higher compared to the regions without CNAs. In addition, the direction of CNAs turned out to be in the direction of SGAs in each stage, which is in the line with the notion that CNA is one of the mechanisms of somatic gene deregulation^27^. However, the direction of IDC-CNAs was in the opposite direction of SGAs inherited from earlier stages. We also found that the function of SGAs inside and outside of CNAs could be formulated in a *forward evolution* model with additive impact on the fine-tuning of operating signaling pathways of tumors. Surprisingly, we found that CNAs work partly in a *backward evolution* model where this mechanism removes the negative regulators of selected signaling pathways during tumor progression. A recent punctuation evolution model speculated a driving role for CNAs as they found the majority of CNAs are acquired at the earliest stages of tumor evolution followed by stable clonal expansions forming the tumor mass^28,29^. Our analyses suggest that the evolution of tumors and especially the malignancy trait of tumors are mostly related to earlier SGA events out of CNAs as they were the main components of perturbed pathways in IDC. In addition, the selected CNAs in later stages added no new functional pathway for the operating signaling pathways of advanced tumors as compared to DCIS. Moreover, the *backward evolution* mechanism that was detected in the CNAs leaves hardly room to interpret CNAs as the main driving force of tumor evolution but rather as fine tuner for the function of SGAs.

We further found that Met samples have equal similarities to DCIS and IDC for the shared perturbed signaling pathways and driver genes. However, Met has more unique disturbed pathways compared to DCIS and IDC. This probably reflects a higher chance of Met samples for the neutral evolution as this is shown to be more prominent in the absence of selection pressure codition^18,26^. Therefore, we speculated an early divergence of metastases at DCIS time point that we corroborated in our phylogenetic analyses. This is in the line with recent studies that found metastatic dissemination of breast and colon cancer cells is higher in earlier stages of tumor progression compared to the advanced tumors, and these early-disseminated cancer cells are precursors of a great proportion of metastases^10,11,30^. In addition, recent evidences indicated the uniformity of driver gene alterations, even at the single nucleotide level in primary tumor samples and metastases of breast as well as pancreatic cancers^31-33^, suggesting an early divergence of metastases from primary tumor. Moreover, spatial analyses of invasive foci at the DCIS sites showed that genomic evolution occurred prior to the invasion through a multi-clonal invasion model^34^.

## Conclusion

Altogether, this study suggests that the differences in the gene expression profiles of pre and post invasive breast cancer do not present a new founder route of malignancy. Indeed, fundamental genetic alterations are SGAs occurring in the earlier stage of tumor development and further SGAs in later stages, in a *forward evolution* model, serve as a fine-tuning mechanism of those primary changes. Remarkably, we found a *backward evolution* model for the cancer genome evolution which CNAs are selective events and serve as means to reverse expression/direction of a proportion of earlier SGAs. Our methodology introduces a new pipeline for cancer evolution studies and our results draw the global portrait of malignancy evolution of breast cancer.

## Supplementary figures

**Figure S1.**
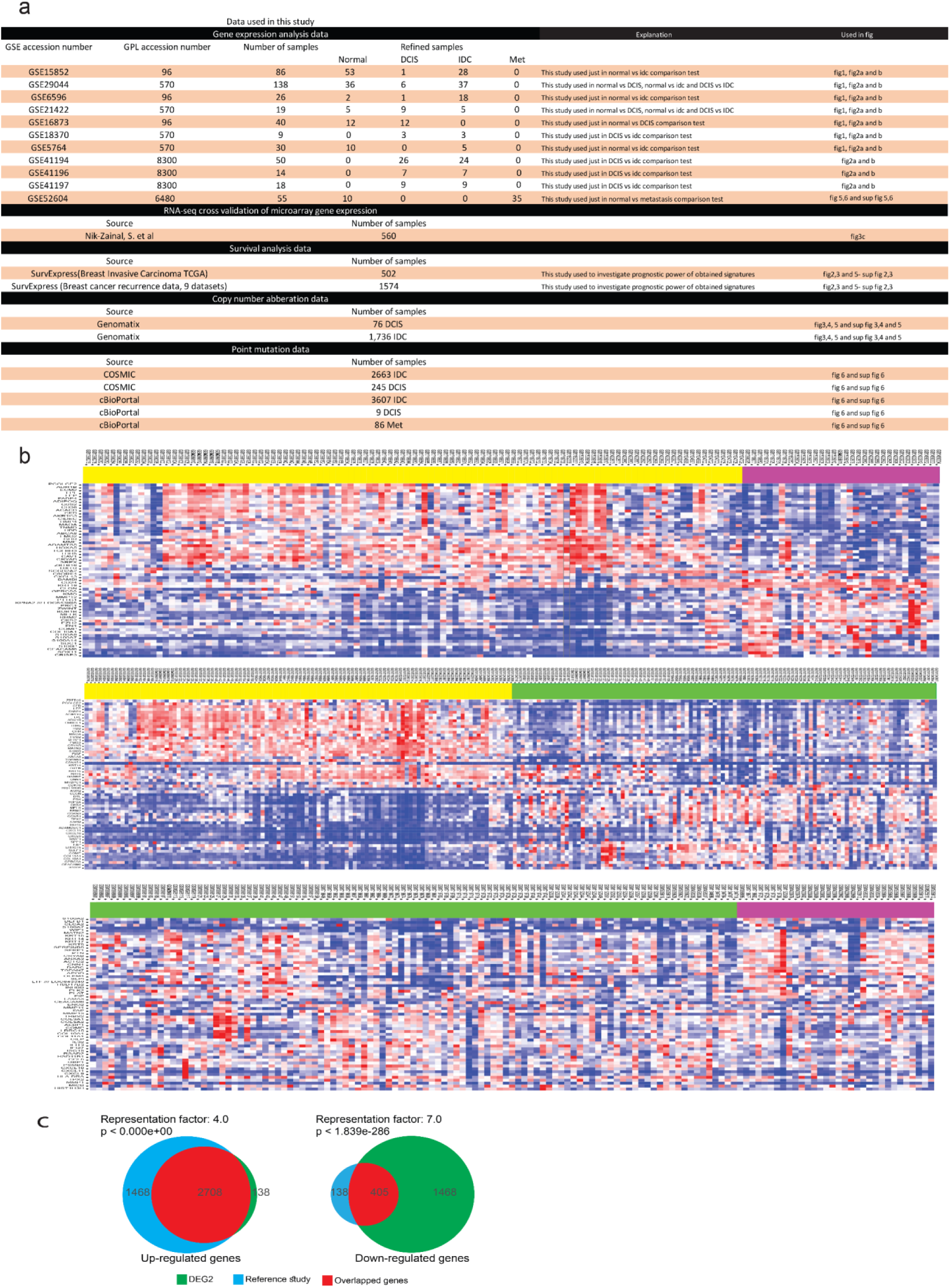
Samples and gene expression heatmaps. **a**) List of samples, datasets and their characteristics used in this study. **b**) Heatmap of 60 genes with the highest LFC in Normal, DCIS and IDC samples. Rows (genes) clustered by Pearson correlation method and average-linkage hierarchical clustering method and samples (columns) clustered by their phenotypes (Hierarchical clustering was not applied on columns). **c**) Venn diagrams show high overlap between DEG2 (IDC vs Normal) and analogous RNA-seq data^22^. The *p*-value calculated by the exact hypergeometric probability test. The RF that presented the number of shared genes divided by the expected number of shared genes resulting from two independent groups. RF more than 1 shows more overlap than expected and vice versa.

**Figure S2.**
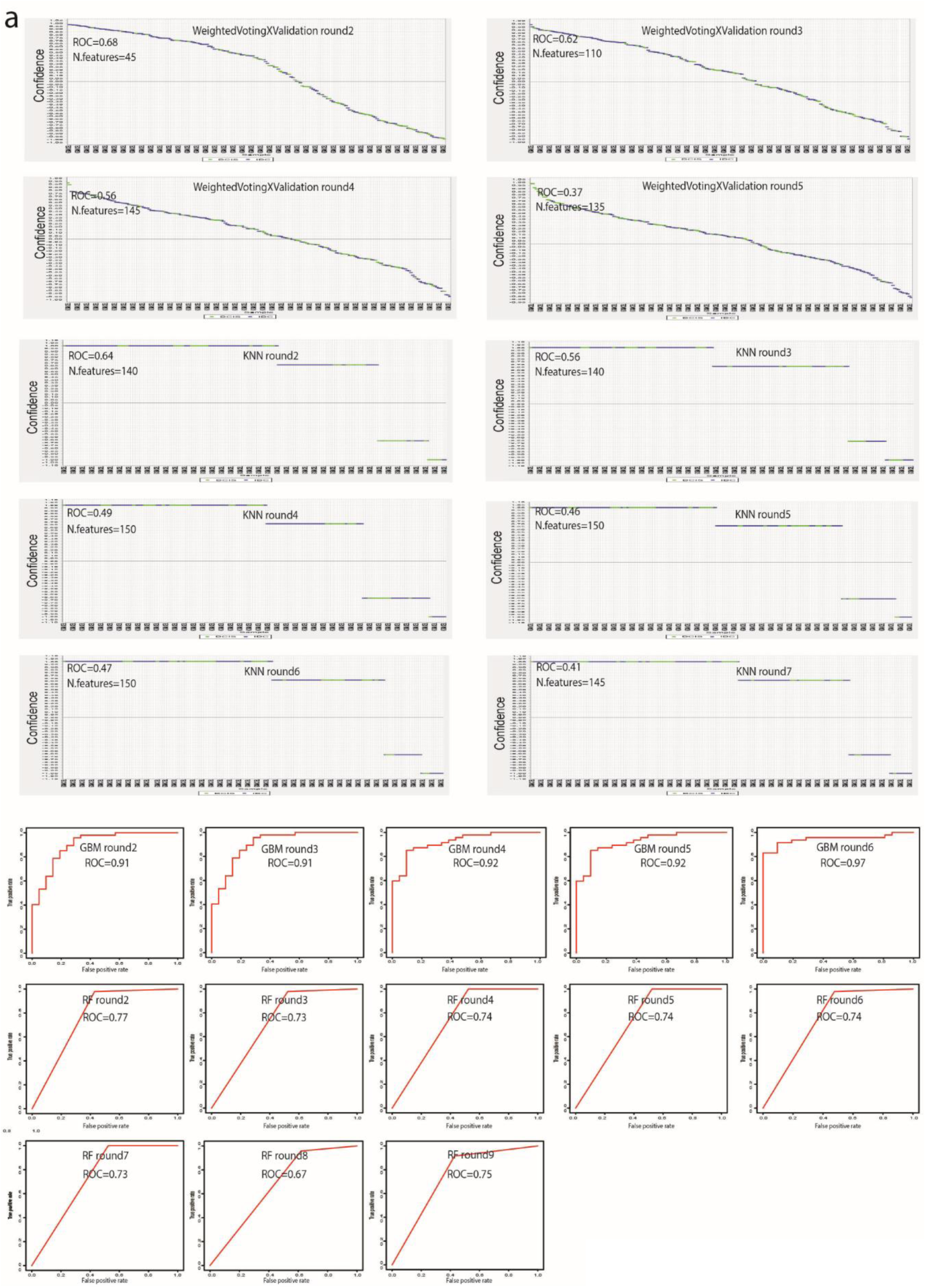
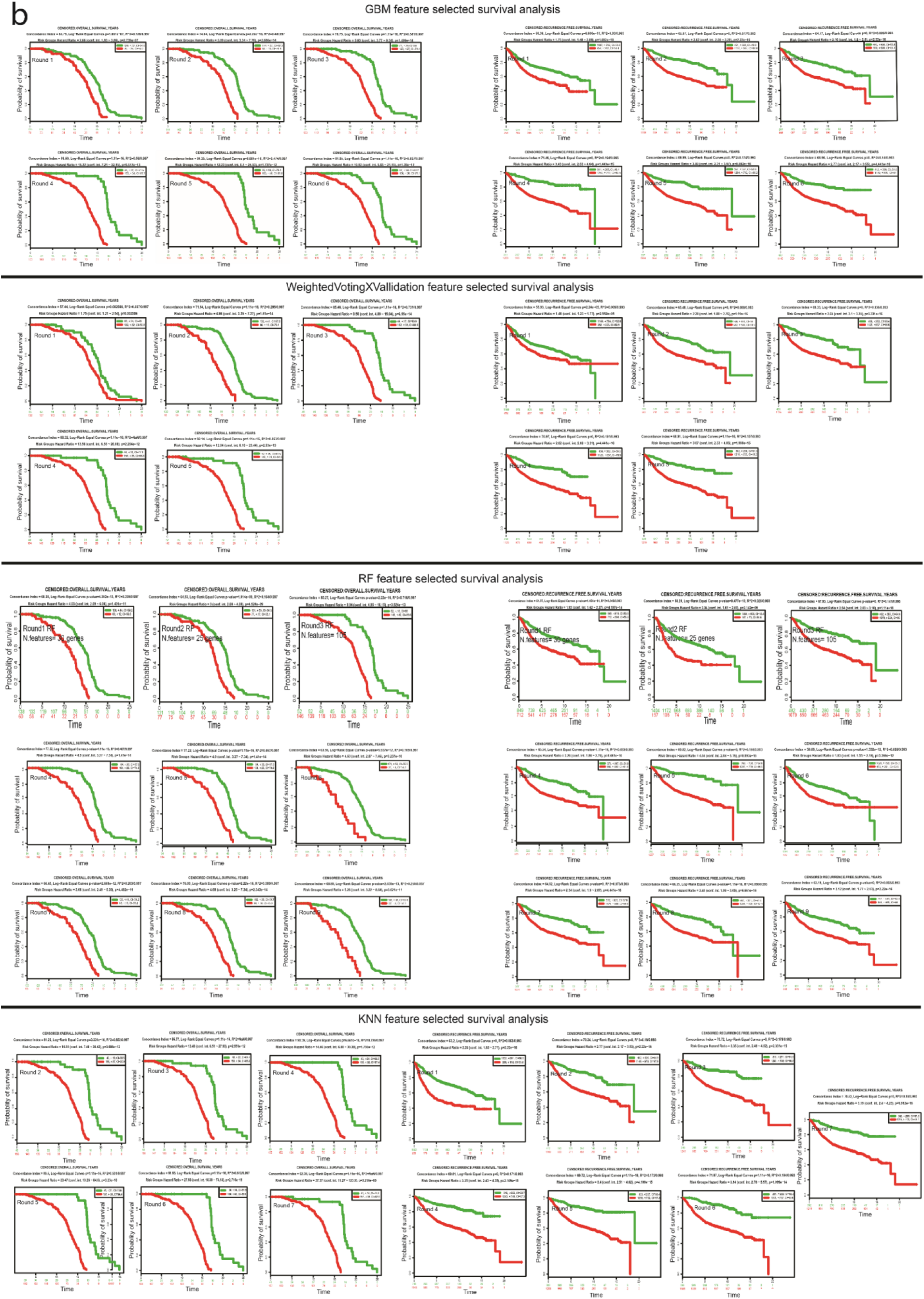
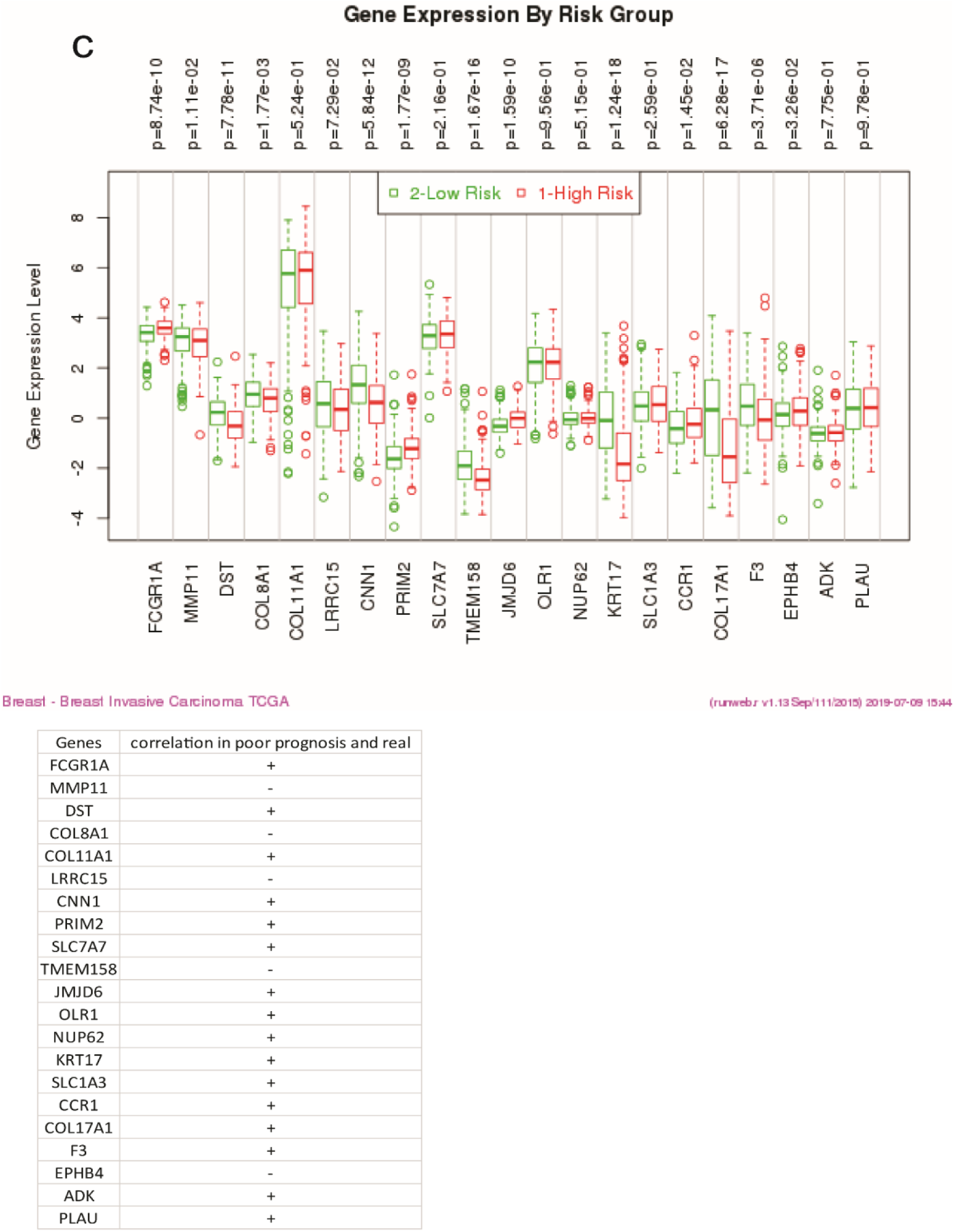
Prediction of the results of machine learning models and survival analyses. **a)** Machine learning (ML) class prediction models that were optimized by four different approaches including GBM, RF, KNN and weightedVotingXValidation. Plots represent specificity and sensitivity of models through ROC values for the features, which are shown in Fig 2b. **b**) Kaplan-Meier survival analyses (OS and RFS) for gene features selected in each round of ML models presented in Fig 2b. **c**) Example of correlation between gene expression in the poor (red) and good (green) prognosis samples obtained for survival analyses. Table in the below of gene expression graph represents the concordance between expression of that given gene in poor prognosis samples and the real expression direction in the IDC samples.

**Figure S3.**
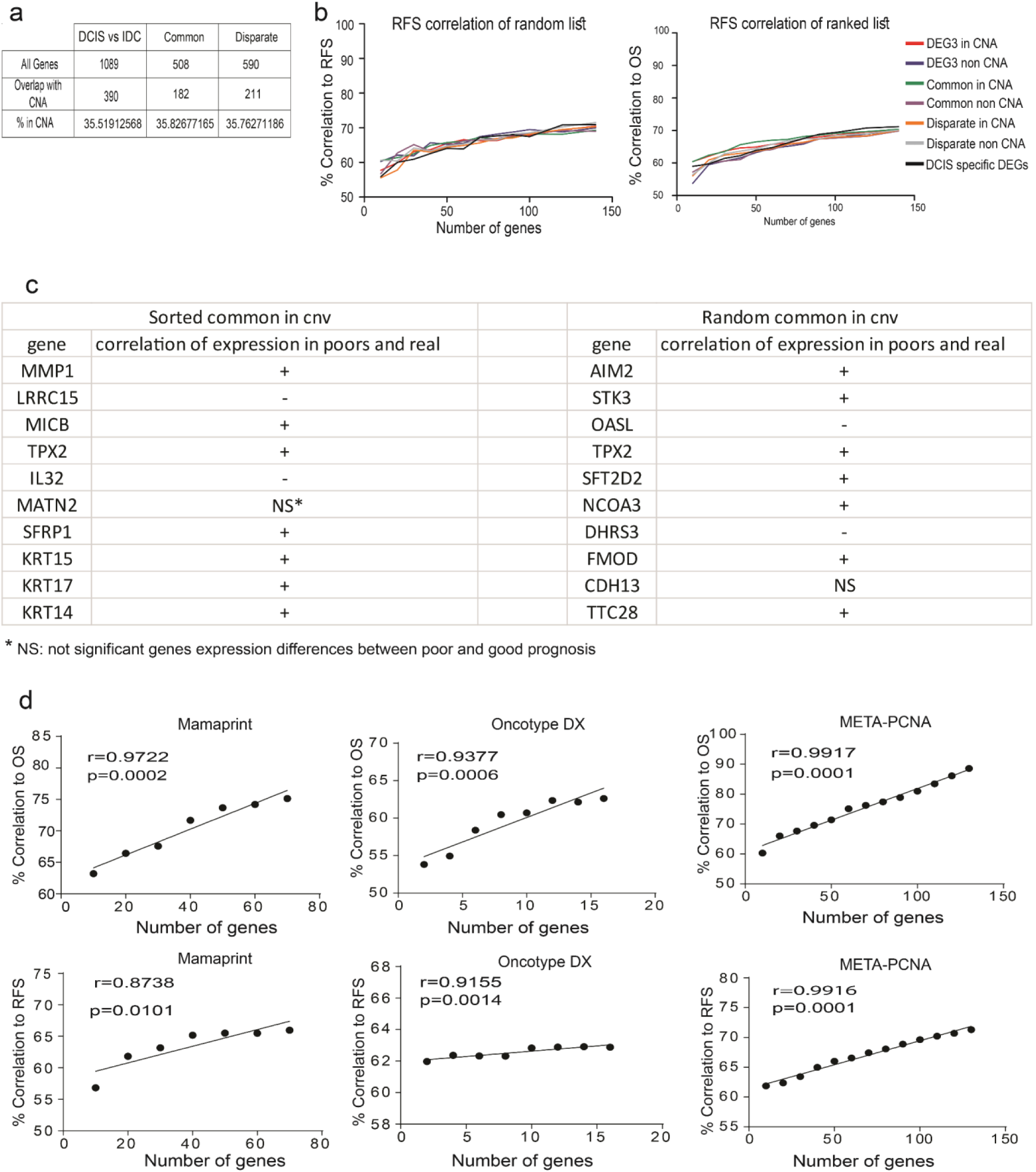
Correlation of number of genes and survival. **a**) Number of common and disparate genes and their contribution in CNAs. **b**) Correlation of number of genes selected randomly (left) or based on ranked fold changes (right) to relapse free survival. Genes are selected from common, disparate, DCIS specific and DEG 3 (IDC vs DCIS) within and out of CNV. **c**) Two example of correlation between gene expression in the poor and good prognosis samples obtained for survival analyses in Figure 3d. Table represents the concordance between the expression of that given gene in poor prognosis samples and the real expression direction in the IDC samples. **d**) Correlation of CI of overall survival and recurrence free survival CI with the number of genes from available gene signatures including Mamaprint, META-PCNA and Oncotype DX. The *p*-values in panel d calculated by Pearson correlation test.

**Figure S4.**
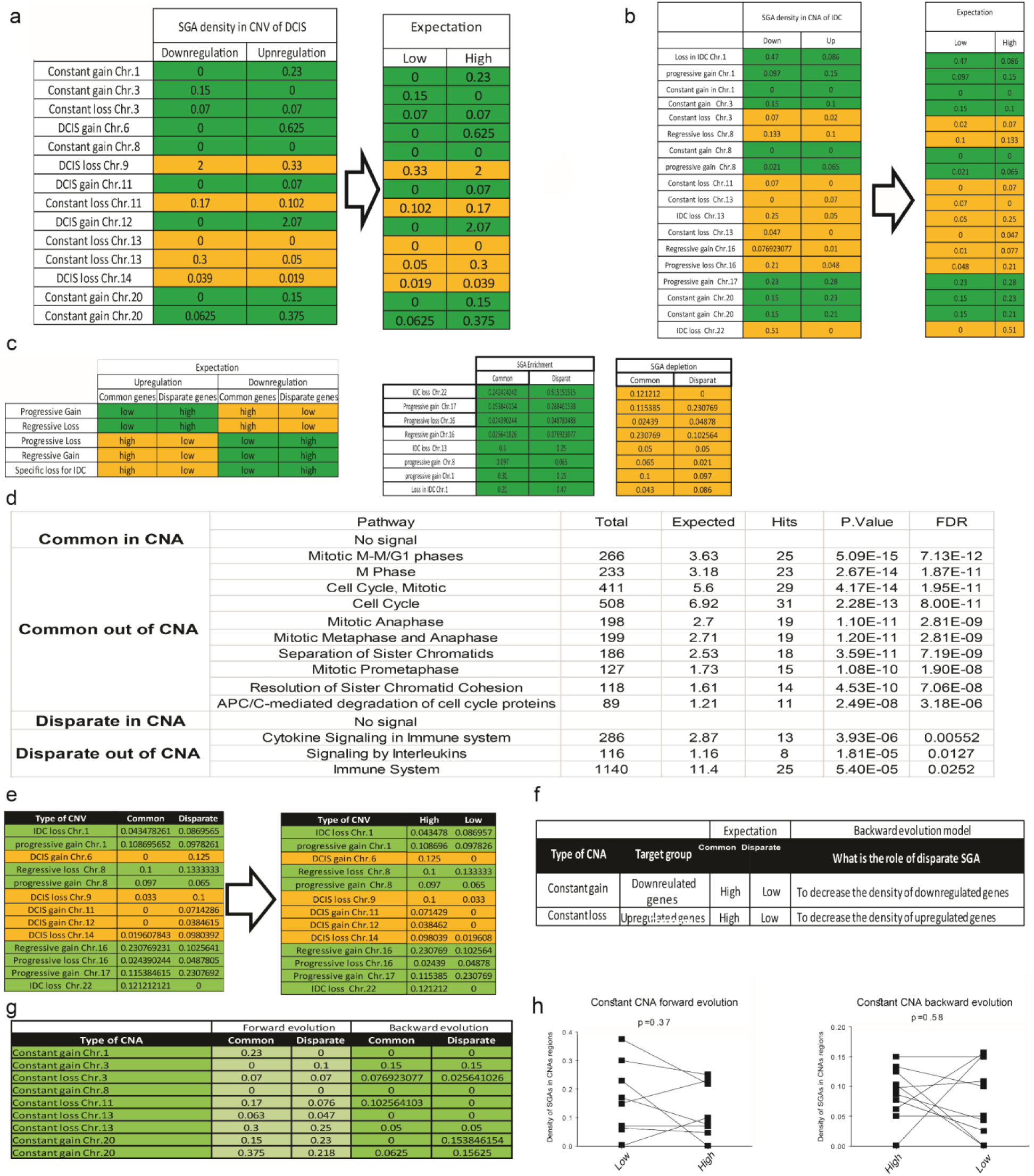
Direction of changes in CNA and SGA. **a** and **b**) Scheme table of the density of SGAs (number of SGAs per Mb) in DCIS-CNA (a) IDC-CNA (b) prototypes show in the left table and the direction of our predictions show in the right table. The yellow squares show loss-CNAs where we expect the density of down-regulated genes would be high. Therefore, in the right table we flipped the numbers of yellow squares to bring “down” numbers in the “high” columns. The green squares present unchanged numbers between left and right columns. **c**) Scheme tables present *forward evolution* model for relation of SGAs through DCIS (common SGAs) to IDC (disparate SGAs). Note that we evaluated all SGAs (common and disparate) in the IDC-CNAs prototypes as the final evolved version of CNAs. The green color predicts the enrichment of SGAs and the yellow one predicts the depletion of SGAs. The right tables present real numbers of SGAs density. **d**) Enrichment pathway analysis of common SGAs within CNAs (CWC), common out of CNAs (COC), disparate within CNAs (DWC) and disparate out of CNAs (DOC) networks by Reactome enriched functional subnetworks in the main PPIN. **e**) This table presents the density of SGA numbers for CNA-prototypes with a possible *backward evolution* function which is mentioned in the table in Fig 4i. **f**) Scheme table for the *backward evolution* concept in CNA-constant prototypes. **g**) SGAs number for *forward* and *backward evolution* models in the CNA-constant prototypes **h**) Evaluation of *backward* and *forward evolution* model in constant CNAs. *p*-values in panel (h) calculated by Wilcoxon matched-pair signed rank test.

**Figure S5.**
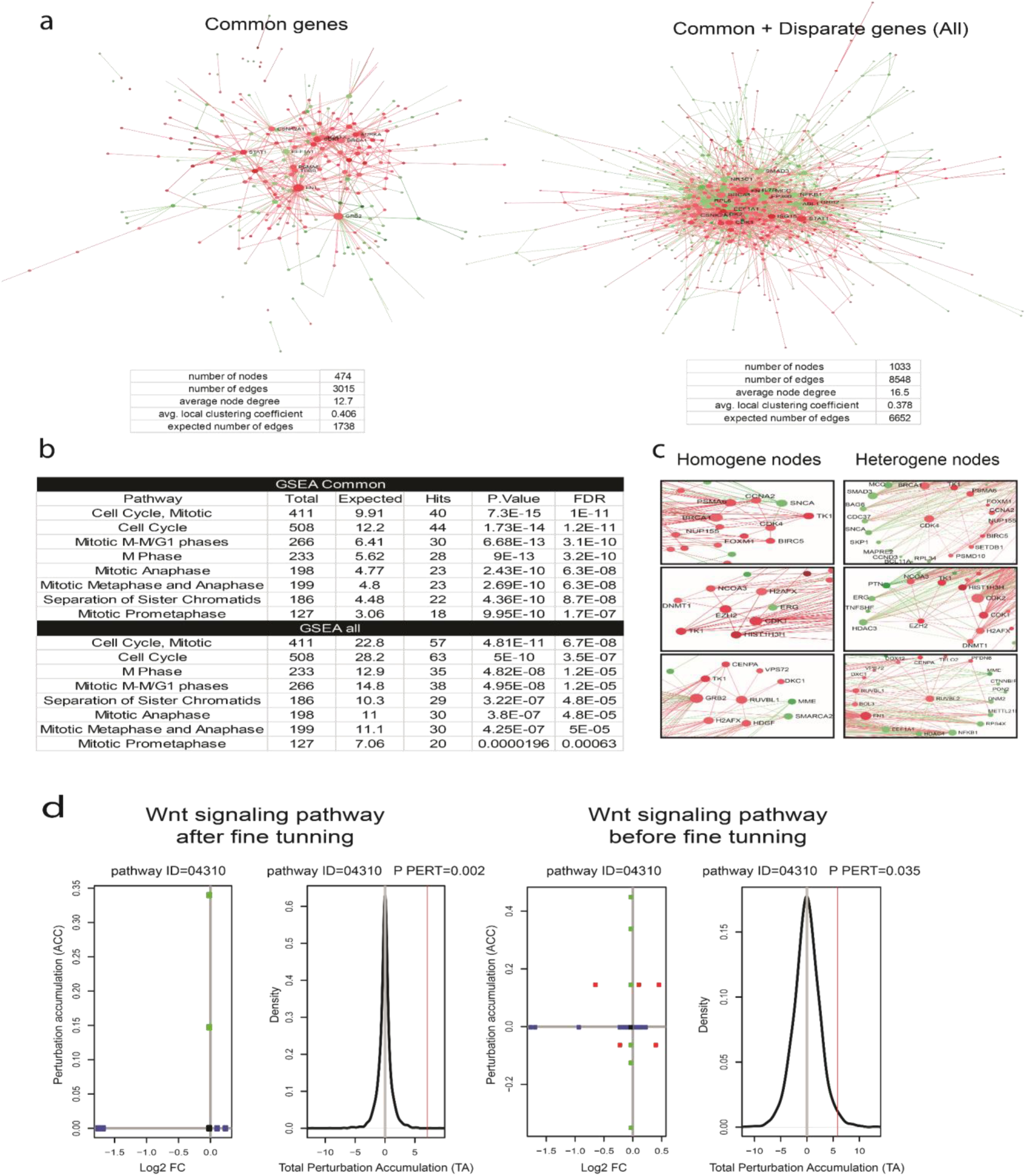
*Forward* and *backward evolution* and *fine-tuning* concepts. **a**) PPINs and its related-statistics associated to common SGAs and all DEG3 (all common and disparate SGAs). PPINs were created by mapping gene sets to the Networkanalyst database. **b**) Enrichment pathway analysis for common and all DEG3 networks (in panel a). Top 10 enriched pathways presented for each gene set profile. **c**) Examples of homogene and heterogene nodes. **d**) Examples of SPIA analyses for DEG3 before and after adding eight *backward evolution* genes. In this plot, the horizontal axis represents the *p*-value (minus log of) corresponding to the probability of obtaining at least the observed number of genes (NDE) in the given pathway randomly. The vertical axis represents the *p*-value (minus log of) corresponding to the probability of obtaining the observed total accumulation (tA) or more extreme on the given pathway randomly. Unchanged genes are assigned 0 log2 fold-change. The null distribution of the net accumulated perturbations is also given (right panel). The observed net accumulation tA with the real data is shown as a red vertical line (see Ref 19 in the main text).

**Figure S6.**
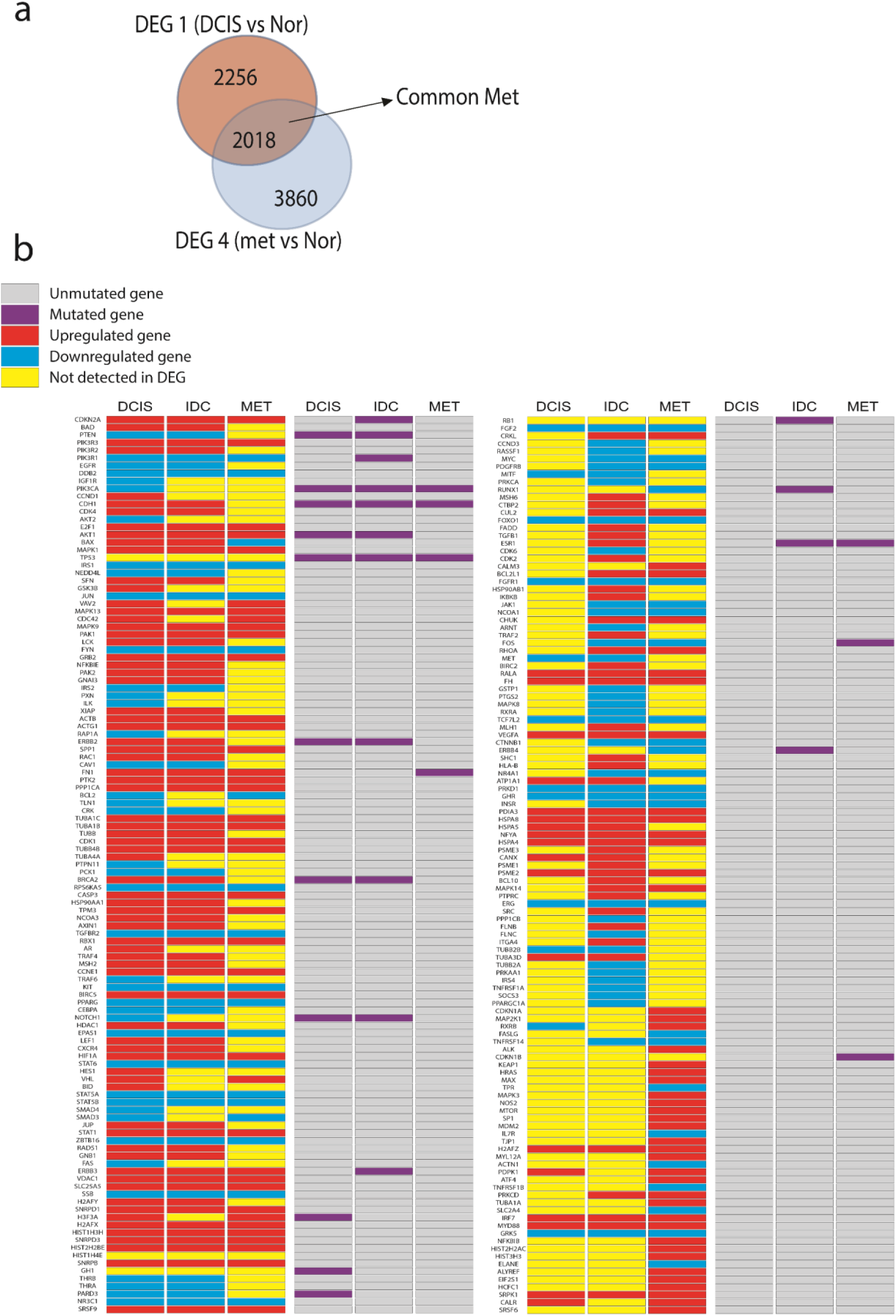
Common-Met and heatmap of driver genes. **a**) Graphic demonstrates common-Met gene expression profile (SI File 8). **b**) Heatmap of all driver genes in DCIS, IDC, and Met with the direction of expression and mutation state.

## Materials and methods

### Data collection, preprocessing, and identification of differentially expressed genes

Gene expression microarray raw data were retrieved from GEO database (see GEO accession numbers in Fig S1a). Then, we discarded low-quality samples (102 samples were excluded) to focus on changes that only trigger the transition between normal to DCIS and DCIS to IDC or metastasis without extraneous interferences. All samples with no clear clinical annotation were also removed. Afterwards, Affy batch objects were created by the affy package^35^ in R and the GCRMA package^36^ was used for preprocessing the datasets (including background correction, normalization, log transformation, and quality control). The identification of differentially expressed genes (DEG) was done in four states including normal *vs* DCIS, normal *vs* IDC, DCIS vs IDC, as well as primary *vs* metastasis via limma^37^ through NetworkAnalyst^38^. The combined *p*-values for the DEGs identification have been obtained by using the state combined Fisher’s method (Adjusted *p*-values for DEGs from each study combined by Fisher’s method). DEGs within the mentioned states are named DEG1, DEG2, DEG3, and DEG4 for normal *vs* DCIS, normal *vs* IDC, DCIS *vs* IDC states, and primary *vs* metastasis respectively (see all DEGs in SI File 1). In order to evaluate our results obtained by microarray technology with RNA-seq data, the DEG2 signature compared with the analogous DEG list from the largest available RNA-sequencing dataset in breast cancer and calculated the significance by exact hypergeometric probability^22^.

### (Un)supervised clustering and machine learning

In an independent analysis, all samples obtained from the GPL96 (HG-U133A) and GPL570 (HG-U133_Plus_2) platforms, were preprocessed by fRMA package ^39^ and merged by COMBAT method ^40^ to remove their batch to batch variations and minimize the study bias effect. The reason why the FRMA preprocessing and COMBAT methods were used for the unsupervised clustering section is that the FRMA is a cutting-edge method for preprocessing when gene expression studies supposed to be integrated directly. Also, COMBAT was used for removing batch effect between studies and constitutes the meta-dataset to perform a reliable visual inspection of relation between the phenotypes. The hierarchical clustering of the 60 most reproducible genes (genes with highest log fold change values and *q*-value<0.01) in the meta-dataset (which was created after merging the individual GSE datasets) was done by the pairwise average-linkage method^41^. Furthermore, PCA was performed in order to identify the correlation among biologically distinct samples. In the supervised class prediction section, all GCRMA normalized data was merged by COMBAT and fed into the Machine Learning (ML) models. In this study, four different approaches of supervised ML for class prediction were used to properly assign samples to DCIS and IDC classes. All data was fed into the models to develop the predictive models by WeightedVotingXValidation^42^ and k-nearest neighbor (KNN)^43^, two modules that ran through Genepattern (http://software.broadinstitute.org/cancer/software/genepattern/)- which use Leave-One-Out Cross-Validation approach to optimize the predictive models, but for Gradient Boosting Machine (GBM)^44^ and Random Forest ^45^ models, the data was split randomly and assigned to train and test datasets. All models obtained high-resolution discriminative features and were optimized by the minimum number of genes and maximum receiver operating characteristic area under curve (ROC AUC). Finally, we developed a pipeline to iteratively evaluate the discriminative genes extracted from each ML model. We excluded top-ranked predicted genes and re-ran models for maximum nine rounds to prioritize each phenotype specific features and to test their correlation with the survival of patients.

### Survival analysis

The prognostic performance of gene signatures was assessed via the cox model in terms of Overall Survival (OS) and Recurrence Free Survival analysis (RFS) using the Survexpress dataset^46^ including 1,574 samples. The survival analysis was applied to highlight whether discriminative genes of each round of ML models have a correlation between top-ranked genes obtained by ML models and to optimize the sample separation of DCIS and IDC. We assume that top-ranked discriminative genes between these two phenotypes (DCIS *vs* IDC) must have a higher hazard ratio and correlation according to the Concordance Index (CI is an index that correlate with the reliability of created cox model^46^). Furthermore, the survival analysis in term of survival months was done using breast invasive carcinoma TCGA datasets including 502 invasive samples for the detected genes involved in *backward evolution*. In all survival analyses, the maximize risk groups option was checked in order to optimize the risk group splitting using Survexpress algorithm.

### Integration of copy number alteration and single gene alteration

Copy Number Alterations (CNA) data was obtained from the Progenetix database (https://progenetix.org/, version 2017) including 76 DCIS and 1,736 IDC samples (SI File 2). A prototype map of CNA was generated for chromosomal regions, which had more than 10% gain or loss of alteration. Progressive gain and loss prototypes were defined for regions, which already had a gain or loss in DCIS samples and further increased gain or loss in IDC samples. Regressive gain and loss prototypes were defined for regions which already had a gain or loss in DCIS samples and further decreased gain or loss in IDC samples. In the following, the DEG3 gene list was cross-sectioned with DEG1 and DEG2 (SI File 1) to generate a common and a disparate gene list. DEG3 includes 1,098 genes divided into 508 common and 590 disparate genes lists. Common lists are those genes shared between three DEGs and disparate list are genes shared between DEG2 and DEG3 (SI File 1). Moreover, these common and disparate DEGs lists are divided into Common Within CNA region (CWC; SI File 1), Common Out of CNA regions (COC; SI File 1), Disparate Within CNA regions (DWC; SI File 1) and Disparate Out of CNA (DOC; SI File 1). For the correlation analyses between Single Gene Alterations (SGAs) and CNAs, we calculated the density of SGAs per mega base pair (density of SGA/Mbp) for each CNA prototype (SI File 3).

### Network analysis

Reconstruction, visualization, and statistical analyses of the Protein-Protein Interaction Network (PPIN) were done using NetworkAnalyst^38^ and STRING^47^. PPINs were created based on molecular interactions from multiple biological interaction databases including innateDB ^48^, IntAct^49^, MINT^50^, DIP^51^, BIND^52^, and BioGRID^53^. We reconstructed specific-PPINs for the COC, CWC, DOC, and DWC. Network-based pathways enrichment analysis was performed by Reactome, KEGG and GO modules through NetworkAnalyst. All *p*-values in this section were calculated by the default setting of NetworkAnalyst and STRING.

### Enrichment analysis

Gene enrichment analysis of PPINs were done using Reactome database through NetworkAnalyst. Only significant enriched pathways (corrected *p*-value < 0.05) were considered. Moreover, Signaling Pathway Impact Analysis (SPIA) was applied to assess unusual signaling perturbations in pathways related to common, all (common + disparate) gene sets, and for analyzing fine tuning functionality effect of *backward evolution* related genes. According to SPIA results, KEGG pathways related genes for pathways that significantly perturbed at least in one state of common, all and DCIS specific related pathways (SI File7), retrieved from Molecular Signatures Database (MSigDB, http://software.broadinstitute.org/gsea/msigdb/index.jsp).

### NGS data and point mutation analysis

In order to find mutation signatures related to DCIS, IDC and metastasis stages, we used COSMIC v89, (https://cancer.sanger.ac.uk/cosmic) which contains 2663 and 245 samples for IDC and DCIS respectively. Moreover, 3607 IDC, 9 DCIS and 86 metastatic samples were retrieved from cBioPortal (http://cbioportal.org; see references)^54,55^. Mutation signatures for each cancer stage obtained from the mentioned data by defining 5, 2.5 and 5% as a cut-off frequency for DCIS, IDC and metastasis. We assumed lower cut-off for the IDC samples due to the higher heterogeneity in population.

### Identification of driver genes

We used DawnRank method for detection of driver genes. In principle, this method integrate PPINs with gene expression data and DNA-based somatic alterations that enable us to find the unique driver genes those their changes could deeply perturb the influential pathways^56,57^. On the other word, DawnRank is elevated version of SPIA which is adding genomic data to the analyses. We changed and adapted SPIA for our analyses. Here, we added frequently mutated genes of each stage to the DEGs of that stage and ran PPIN analyses of new gene list. Then we sorted all nodes with the degree ≥20 and ran a KEGG pathway analyses to find enriched pathways for highly connected genes. Finally, cross comparison of our enriched pathways list with the perturbed pathways that characterized with SPIA detected driver genes of each stage.

### Phylogenetic analyses of driver genes

In order to study the evolutionary ancestral relations between different stages of tumor progression, we used mutation map of driver genes of DCIS, IDC and Met, a binary (0,1) data was created which 1 represents genes with point mutation in driver gene lists and 0 shows not mutated genes(see Fig 6k). This binary matrix analyzed by Parsimony analyses module in Past (V 2,17) software by its default setting.

### Statistical analysis

Statistical analyses and estimation of correlations in this study were performed using GraphPad Prism v.6. Correlation significance calculated by Pearson correlation. The *p*-values reported for survival analysis measured by cox regression hazard ratio and log rank tests. All paired tests were performed by Student’s t-test when data points passed from D’Agostino-Pearson normality tests. In case that the variables not distributed normally according to the D’Agostino-Pearson test (*P* ≤ 0.05), we applied the Wilcoxon test. For comparing numbers between different groups we applied Fisher’s exact test or if the sample numbers were at least 5 in each condition the χ 2 tests. All *P* values are two-tailed. All *P* values and statistical tests are mentioned in either figures or legends to test the similarity of gene expression data of microarray and RNA-Seq (Fig S1b) data, we used exact hypergeometric probability test from online tools (http://www.nemates.org/MA/progs/overlap_stats.html).

## Declarations

### Ethics approval and consent to participate

Not applicable

### Consent for publication

Not applicable

### Availability of data and materials

All raw data for presented graphs and statistics are deposited in source data files and all R codes used for analyses will be provided upon request.

### Author contributions

S.S., A.S.Y. and H.H. designed and evaluated core experiments and wrote the manuscript. M.D, N.Z. and O.W. helped with the input in the concept and manuscript.

## Acknowledgment

Not applicable

