## Supplementary figures and images for "Omics integration analyses reveal the early evolution of malignancy in breast cancer"

### sup fig1 .jpg

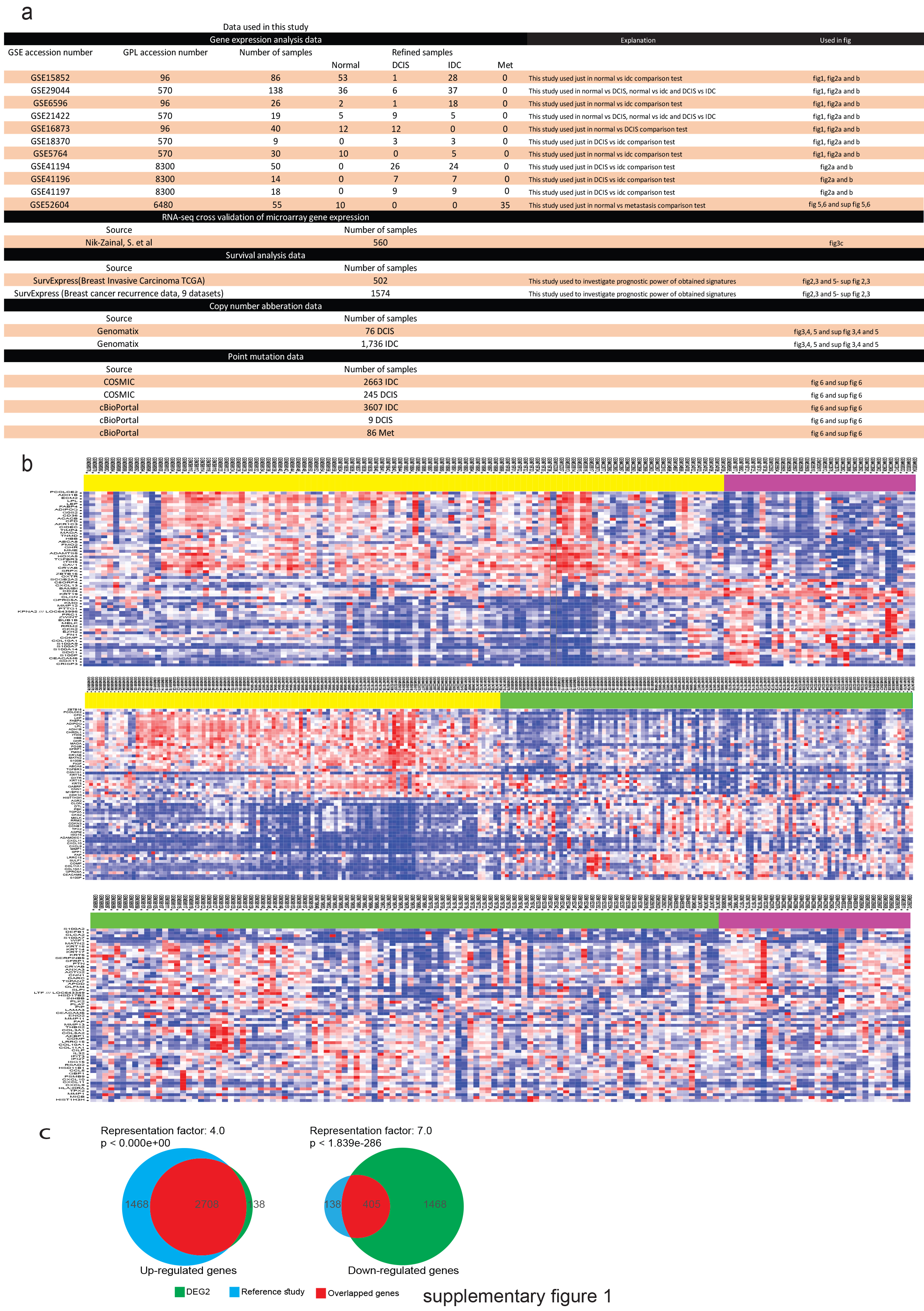

### sup fig2a .jpg

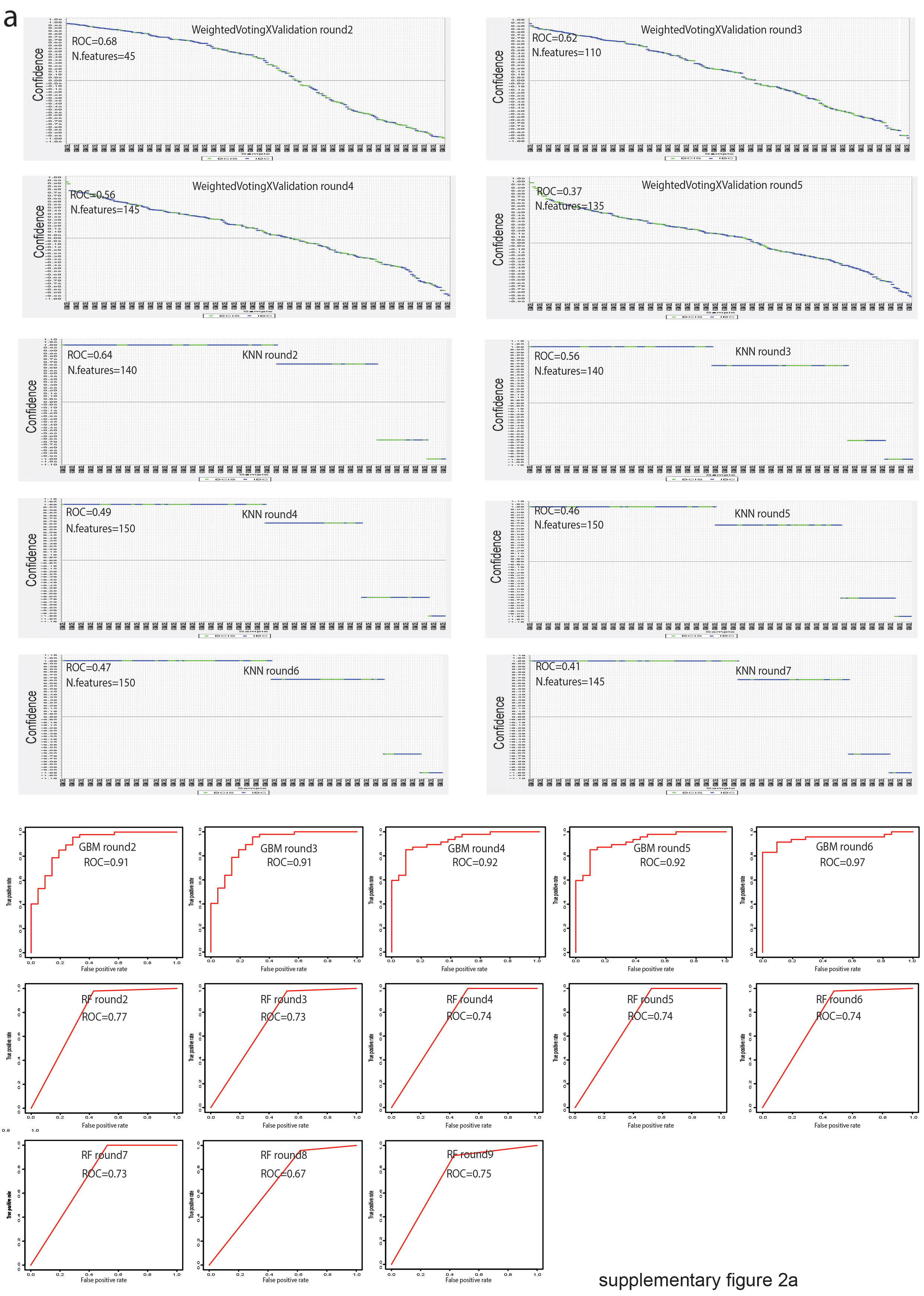

### sup fig2b .jpg

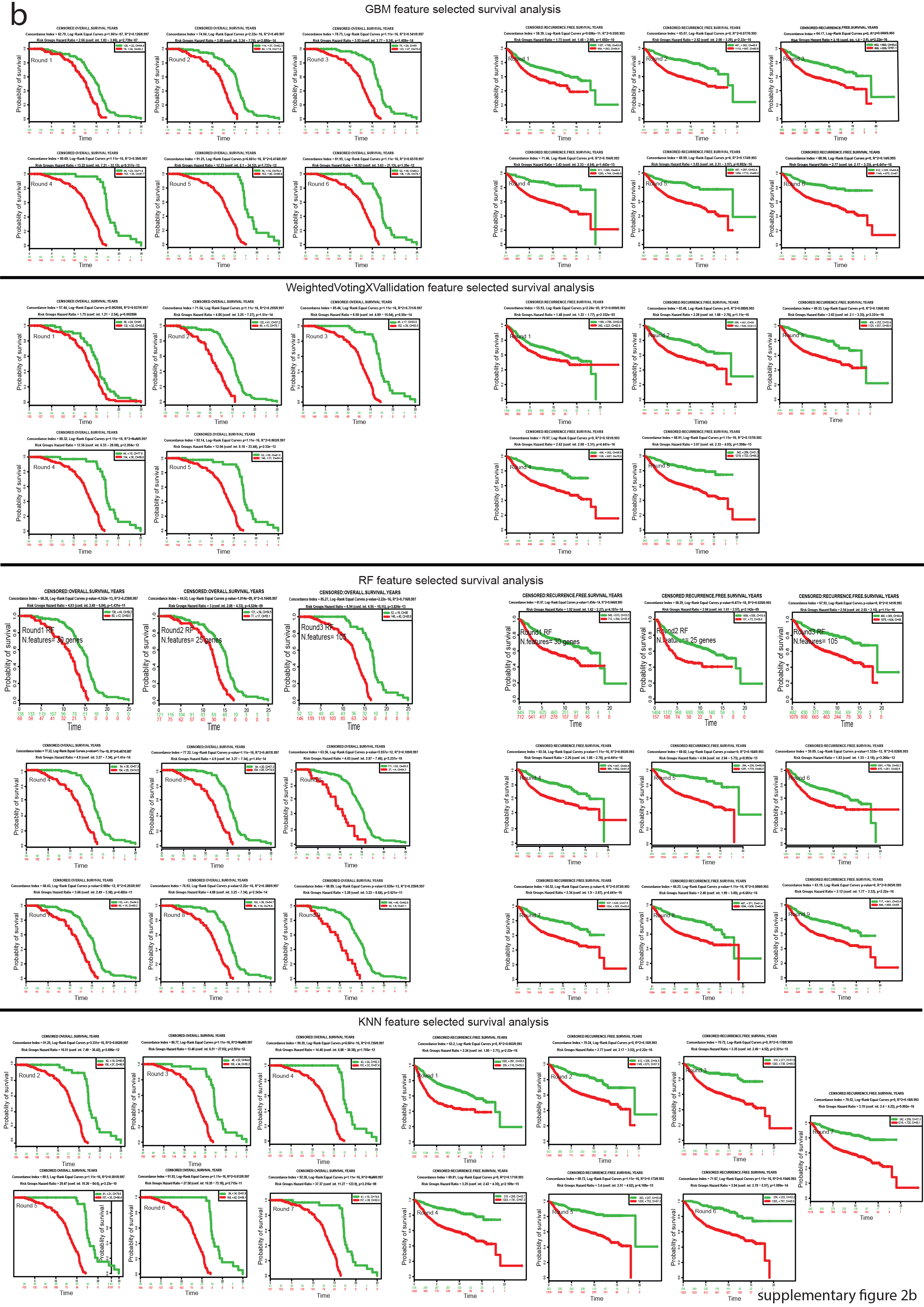

### sup fig2c .jpg

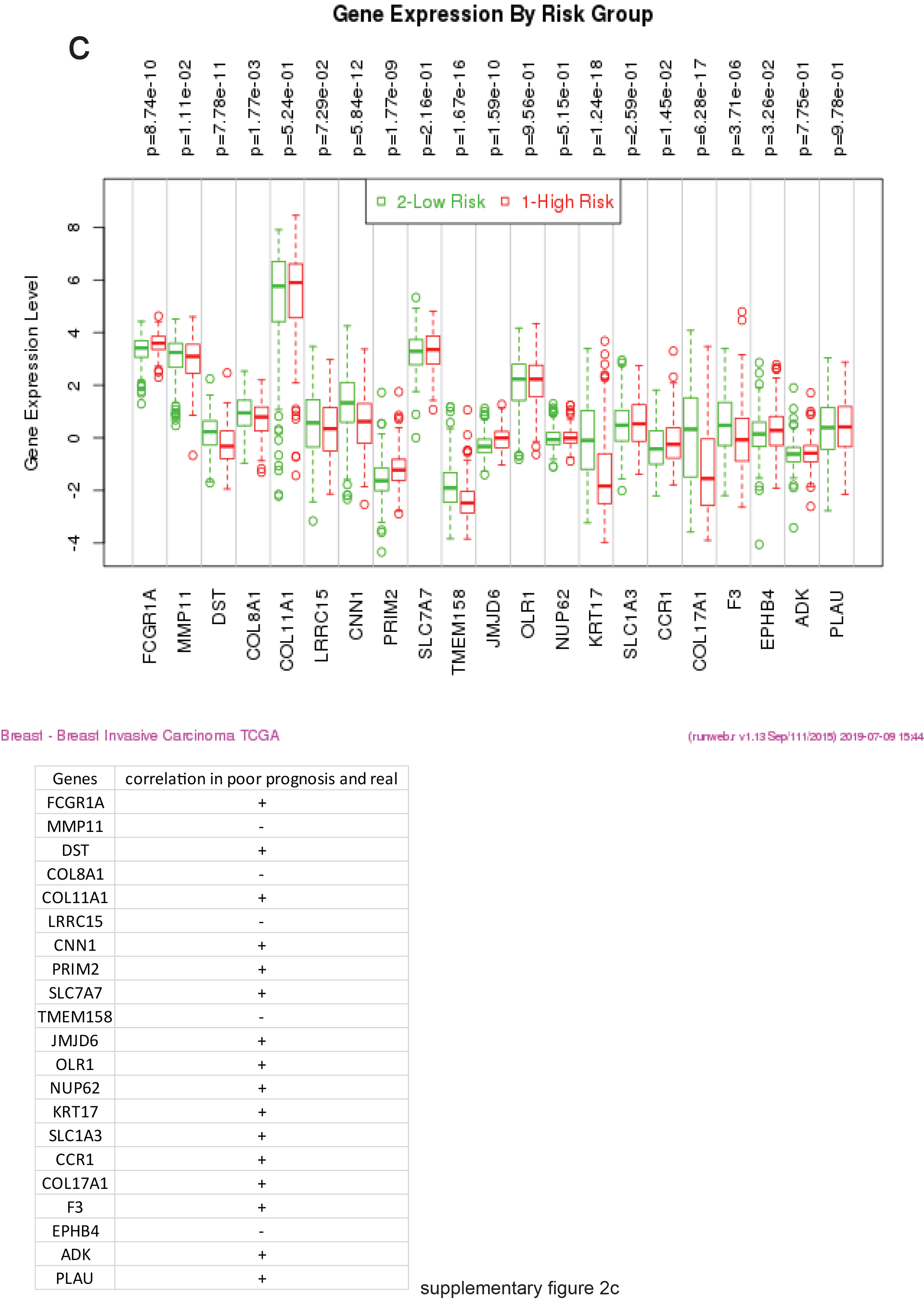

### sup fig3 .jpg

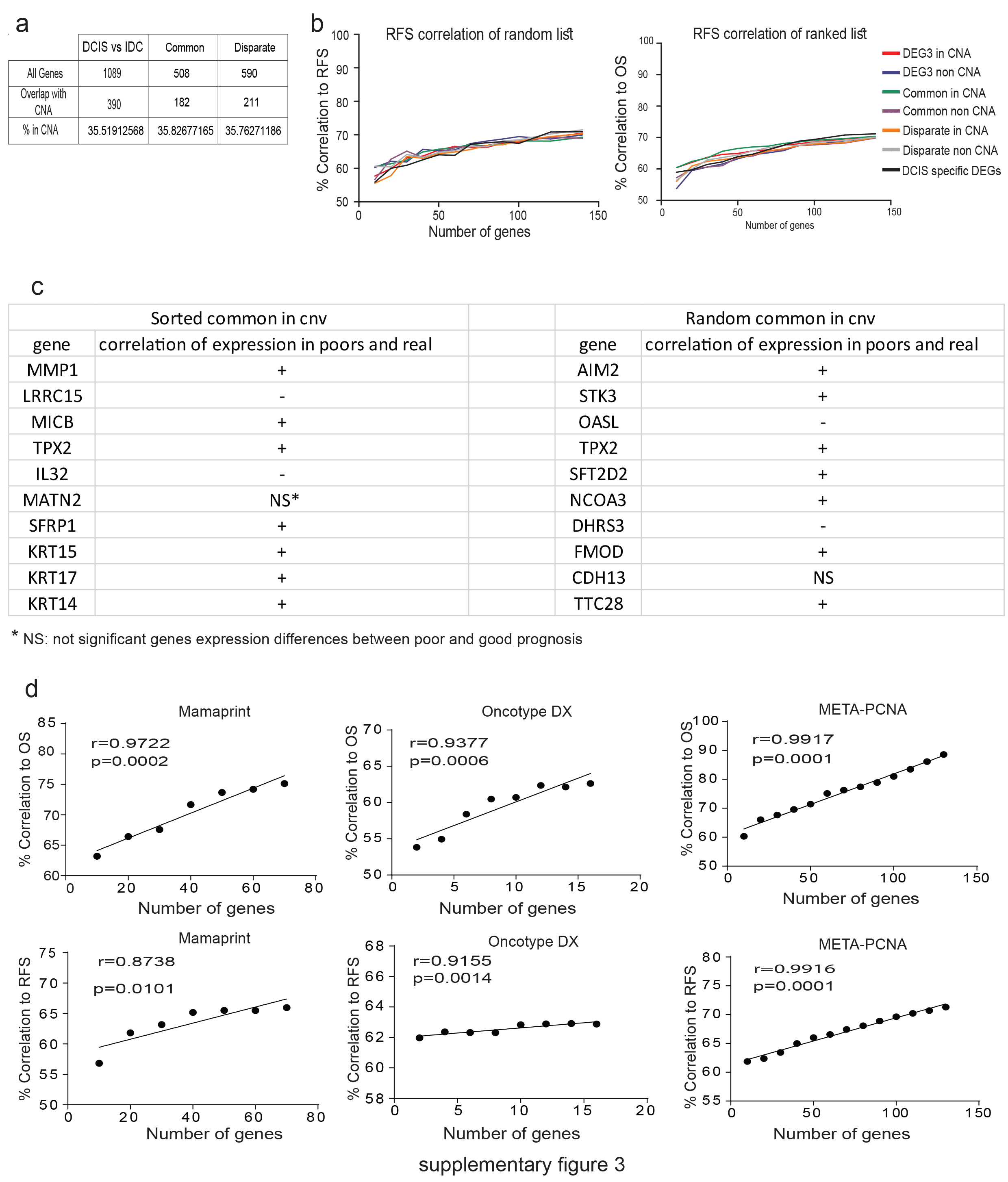

### sup fig4 .jpg

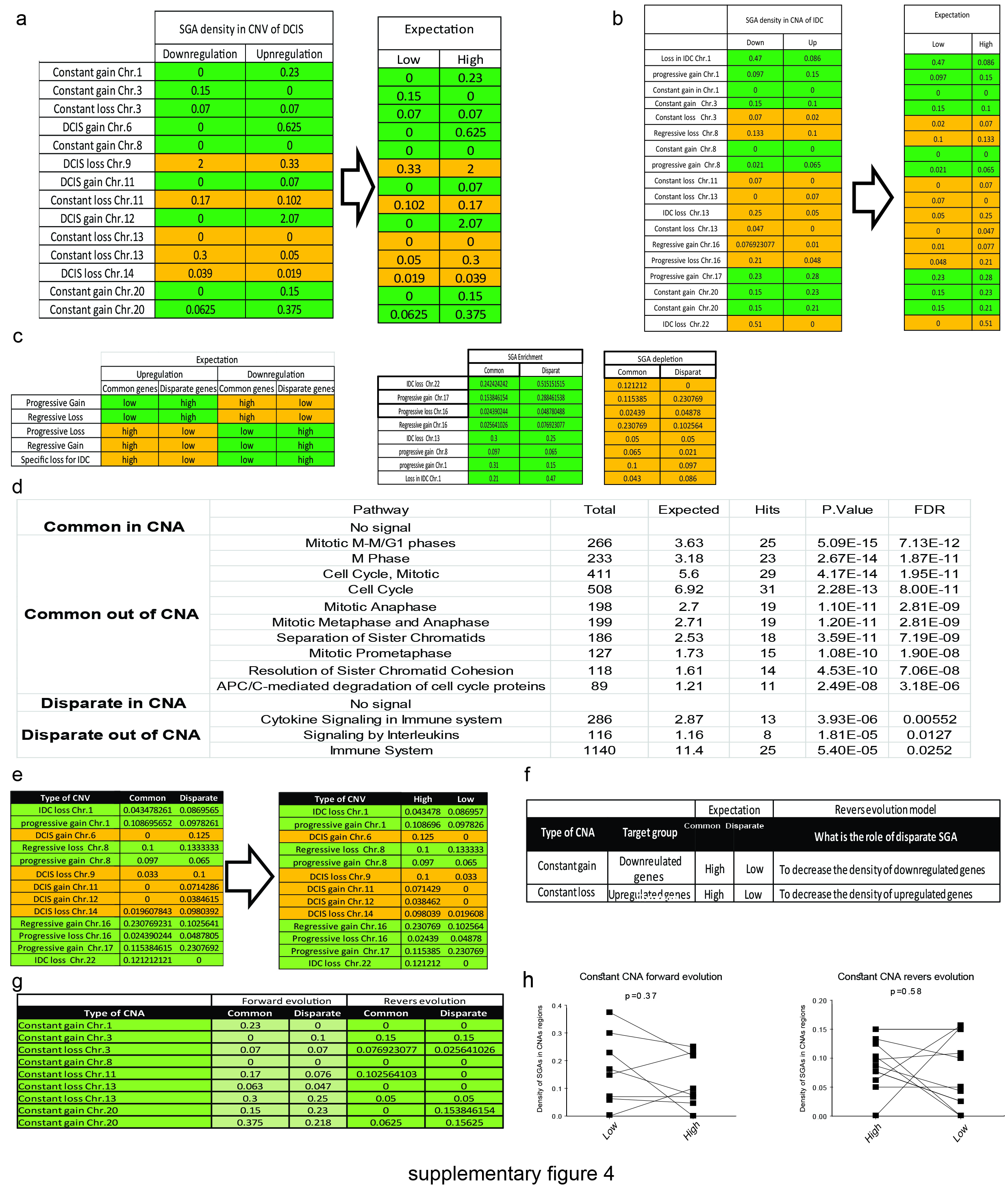

### sup fig5 .jpg

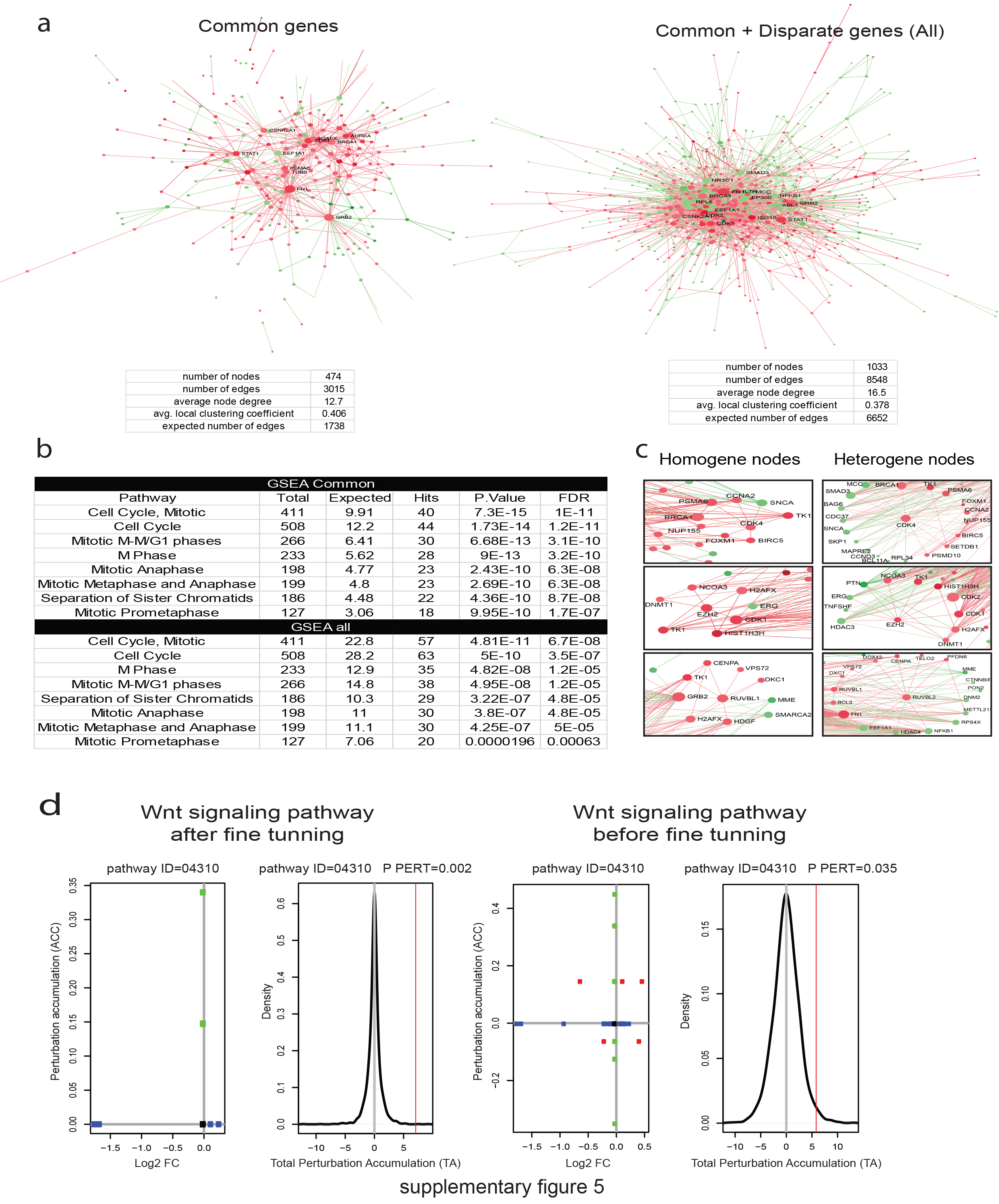

### sup fig6 .jpg

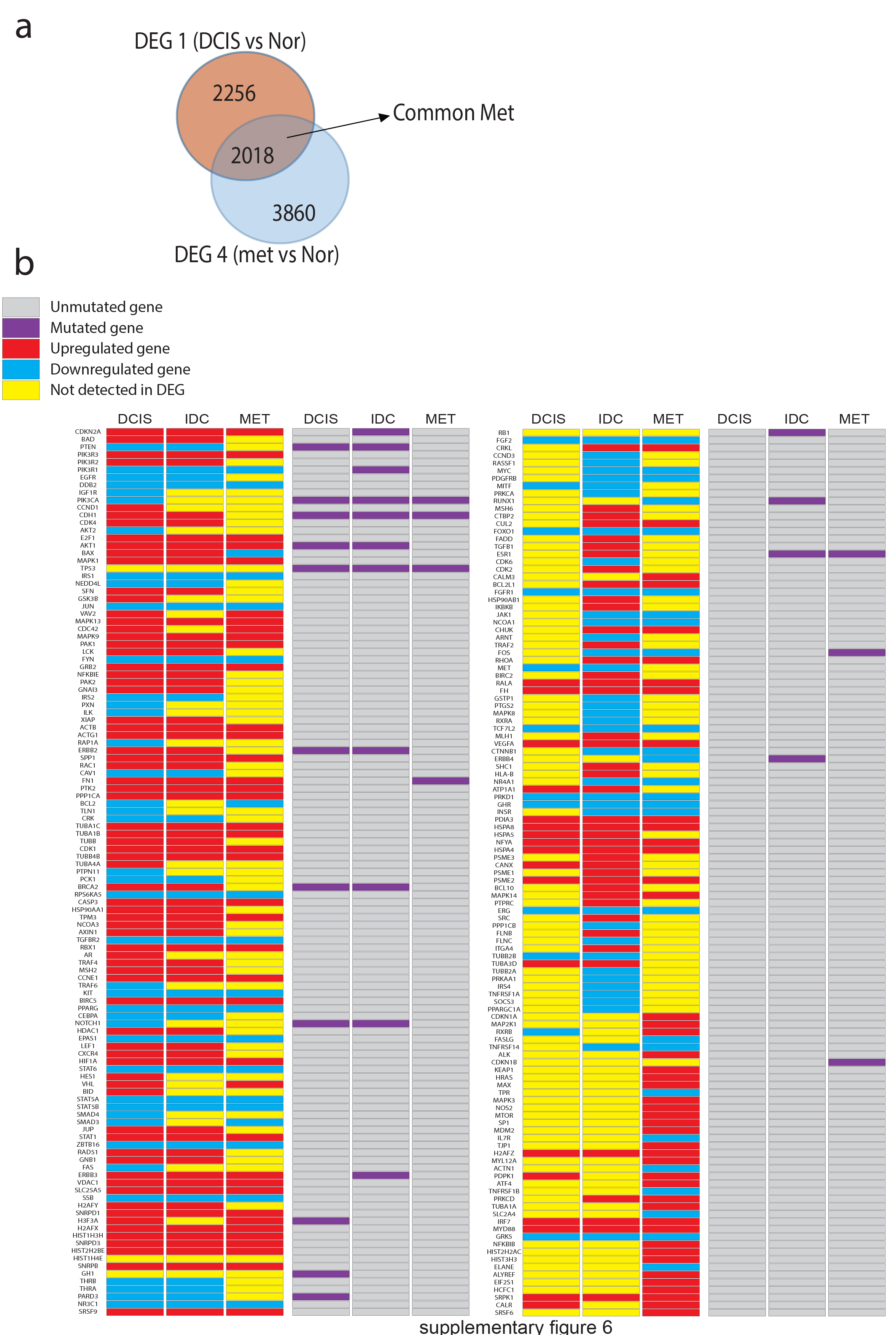
